# Gap junctions deliver malonyl-CoA from soma to germline to support embryogenesis in *Caenorhabditis elegans*

**DOI:** 10.1101/2020.04.24.060160

**Authors:** Todd A. Starich, David Greenstein

## Abstract

Gap junctions are ubiquitous in metazoans and play critical roles in important biological processes, including electrical conduction and development. Yet, only a few defined molecules passing through gap junction channels have been linked to specific functions. We isolated gap junction channel mutants that reduce coupling between the soma and germ cells in the *C. elegans* gonad. We provide evidence that malonyl-CoA, the rate-limiting substrate for fatty acid synthesis (FAS), is produced in the soma and delivered through gap junctions to the germline; there it is used by fatty acid synthase to critically support embryonic development. Separation of malonyl-CoA production from its site of utilization facilitates somatic control of germline development. Additionally, we demonstrate that loss of malonyl-CoA production in the intestine negatively impacts germline development independently of FAS. Our results suggest that metabolic outsourcing of malonyl-CoA may be a strategy by which the soma communicates nutritional status to the germline.

**Impact Statement:** Malonyl-CoA, the rate-limiting substrate for fatty acid synthesis, is produced in the soma and delivered through gap junctions to the germline to promote reproduction and coordinate it with nutritional status.

## Introduction

Gap junctions are clusters of intercellular channels allowing direct passage of small molecules between coupled cells. They are widespread in multicellular organisms, though different gene families encode the protein subunits in invertebrates (innexins) and vertebrates (primarily connexins) (*Skerrett et al., 2017*). While the capability of small molecules to pass through gap junctions can be shown readily in cultured cells, identification of essential molecules that must pass through channels *in vivo* is challenging. Additionally, disruption of gap junction coupling can affect cell adhesion (*Meyer et al., 1992*), and resultant defects in intercellular communication may not necessarily relate to the passage of molecules through junctional channels.

In *C. elegans*, germ cells are coupled throughout their development to the somatic gonad by gap junctions (*Starich et al., 2014*). Germ cells progress in assembly-line fashion through the early stages of meiosis (*Figure 1A*). Gonadal gap junctions have been implicated in germ cell proliferation, oocyte growth, regulation of oocyte meiotic maturation, early embryogenesis and sperm guidance (*Govindan et al., 2006; Whitten et al., 2007; Nadarajan et al., 2009; Edmonds et al., 2011; Starich et al., 2014*). Gap junction hemichannels (half-channels) in the somatic gonad are comprised of the recently duplicated innexins INX-8 and INX-9; apposing germ cell hemichannels are heteromeric and consist of INX-14 in conjunction with either INX-21 or INX-22 (*Figure 1–figure supplement 1*). Germ cells fail to proliferate in *inx-8(0) inx-9(0)* mutants, at a point upstream of, or parallel to, GLP-1/Notch activation of germ cells by Delta class ligands LAG-2 and APX-1. These ligands are produced in the distal tip cell (DTC) that maintains the proliferative stem cell niche (*Austin and Kimble, 1987, 1989; Henderson et al., 1994; Nadarajan et al., 2009*).

**Figure 1.**
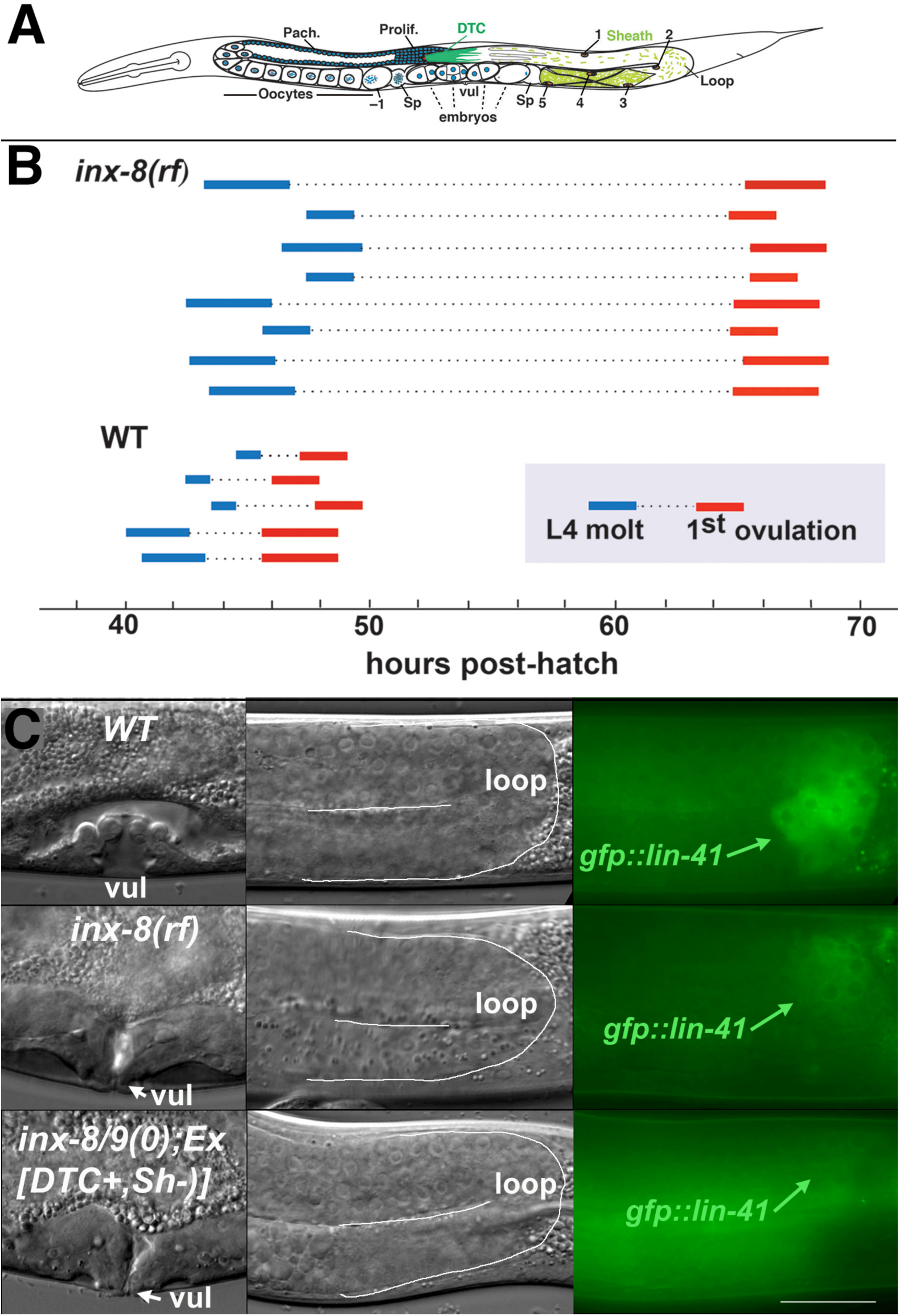
Ovulation and oocyte development are delayed in *inx-8(rf).* (A) Diagram of adult *C. elegans* gonad. Posterior arm shows the positions of the somatic distal tip cell (DTC) and five pairs of sheath cells (nuclei are numbered). Anterior arm shows the underlying germline, approximate regions of germ cell proliferation (Prolif.), meiotic pachytene stage (Pach.), and developing oocytes. Embryos typically undergo some cell division in the uterus before being laid. Sp, spermatheca; vul, vulva. (B) Compared to wild type (N2), ovulation in *inx-8(tn1513 tn1555rf)* mutants (a.k.a. *inx-8[rf]*) is delayed ∼18 hrs. Bars representing 1-3 hr windows indicate approximate time of the fourth larval stage (L4 to adult) molt, followed by time of appearance of the first fertilized embryo ovulated into the uterus. (C) Onset of *gfp::lin-41* expression as a marker of oocyte development is delayed in *inx-8(rf)*. Panels show DIC images of vulval development (left), loop region of gonad arm (middle), and expression of *gfp::lin-41* (right). Wild-type hermaphrodites express *gfp::lin-41* by the approximate L4.7 sub-stage of vulval development (vul); *inx-8(rf)* hermaphrodites do not express *gfp::lin-41* until after vulval development is complete but before the L4-to-adult molt. Null *inx-8(tn1474) inx-9(ok1502)* hermaphrodites, rescued for *inx-8::gfp* expression in the DTC with *Ex[lag-2p::inx-8::gfp]*—a.k.a. *inx-8/9(0); Ex[inx-8(DTC+, Sh-)]—*show a similar delayed onset of *gfp::lin-41* expression as *inx-8(rf)*. Bar, 20 μm.

Germ cell proliferation can be rescued in *inx-8(0) inx-9(0)* mutants to half the level of wild type by driving *inx-8* expression in the DTC with the *lag-2* promoter (*Starich et al., 2014*). In these animals, gap junctions between soma and germline are established in the distal gonad arm, but ostensibly not in the proximal arm; many resultant embryos exhibit characteristics of the Pod phenotype (<u>P</u>olarity and <u>O</u>smotic sensitivity <u>D</u>efect) (*Tagawa et al., 2001*), including eggshell and permeability barrier deficiencies, cytokinesis failures, and polar body extrusion defects. Other genes with mutations producing Pod embryos include those involved in eggshell synthesis, fatty acid synthesis (FAS), the APC/separase pathway, and a coronin-like protein (*Rappleye et al., 1999, 2003; Tagawa et al., 2001; Zhang et al., 2005; Johnston et al., 2006; Olson et al., 2006; Benenati et al., 2009*).

Molecules moving through soma-germ cell gap junctions necessary for proliferation and early embryogenesis are unknown. We use genetic methods to show that malonyl-CoA (mal-CoA) is produced by *pod-2* (acetyl-CoA carboxylase, ACC) in the soma. Transfer of mal-CoA to the germline, for use by *fasn-1* (fatty acid synthase, FASN), is impacted by the integrity of INX-8-containing channels under conditions in which gap junction formation appears normal. Rescue of the Pod phenotype in *pod-2* genetic mosaics indicates a somatic sheath focus of action. Our results support the model that mal-CoA transits from soma to germline through gap junction channels.

## Results

### Gametogenesis is delayed by the *inx-8(tn1513 tn1555)* reduction of function (*rf*) mutation

We previously isolated (in an *inx-9* null background) a strong loss-of-function *inx-8* allele *(tn1513)* with sufficient residual function to contribute to channel formation and cell adhesion but not to support germ cell proliferation (*Starich et al., 2014*). *tn1513* encodes a T239I change in the second extracellular loop of the innexin monomer, predicted to be involved in docking with octameric hemichannels on apposing cells (*Figure 1–figure supplement 1*) (*Oshima et al., 2016*). Using this allele in a genetic screen for restoration of germ cell proliferation, six *inx-8* intragenic suppressor mutations were isolated (to be described elsewhere). One of these, *tn1555*, encodes a D24N change in the amino terminus (*Figure 1–figure supplement 1*), near a site that may associate with a cytoplasmic dome structure surrounding the hemichannel outer pore (*Oshima et al., 2016*). *inx-8(tn1513 tn1555rf)*—for brevity to be referred to as *inx-8(rf)*—partially restored germ cell numbers and fertility to one-third the level of wild type (*Figure 1–figure supplement table 1*). *inx-8(rf)* is rescued by an *inx-8(+)* extrachromosomal array, and rescue of *inx-8(0) inx-9(0)* with an *inx-8(rf)::gfp* array mimics the phenotype of the suppressor mutant; therefore observed phenotypes can be attributed to *inx-8(rf)* and not to other possible background mutations.

**Table 1.** Brood sizes suggest FA synthesis influences embryo output.

| Genotype | Larvae | Dead embryos | Total (embryo output) |
| --- | --- | --- | --- |
| Wild type (N2) | 291±29 (n=33) | 2±2 | 293±29 |
| *inx-8(rf)* | 108±40 (n=90) | 11±6 (n=30) | 119±42 |
| *inx-8/9(0);*<br>*Ex[inx-8(DTC+,Sh-)]* | 2±2 (n=32) | 19±13 | 20±14 |
| *pod-2(0); Ex[pod-2(+)]*<br>high copy (20 ng/μl) | 164±50 (n=31) | 118±41 | 282±59 |
| *pod-2(0); Ex[pod-2(+)]*<br>low copy (4 ng/μl) | 91±24 (n=41) | 156±37 | 247±50 |
| *fasn-1(0); Ex[fasn-1(+)]* | 0.05 (n=80) | 276±18 (n=16) | 276±18 |
| *fasn-1(0); inx-8(rf);*<br>*Ex[fasn-1(+)]* | 17±15 (n=47) | 43±27 | 60±37 |

**Source data 1.** Brood counts for genotypes shown in Table 1.

There is an ∼18 hr delay to the onset of fertilization in *inx-8(rf)* compared to wild type, as determined by relating the timing of first ovulation to the L4 molt (L4-to-adult transition) (*Figure 1B*). This delay correlates with late expression of *gfp::lin41* in relation to L4 sub-stages of vulval development (*Mok et al., 2015*); LIN-41 is the earliest known marker of oogenesis (*Spike et al., 2014*). In wild type at 20°C, *gfp::lin41* appears by sub-stage L4.7 (*Figure 1C*); in *inx-8(rf), gfp::lin41* is expressed after completion of vulval morphogenesis but prior to the L4 molt (*Figure 1C; Figure 1–figure supplement table 2*). Spermatogenesis is also delayed—wild-type sperm appear as early as the L4.7 stage, but *inx-8(rf)* sperm first appear after completion of vulval morphogenesis (*Figure 1–figure supplement table 2*). Once started, egg-laying in *inx-8(rf)* continues apace such that ∼99% of progeny are laid within a 96-hour window (3144/3179 larvae; n=25), similar to wild type (5847/5929 larvae; n=21).

### *inx-8(rf)* gap junction formation is more severely affected in proximal gonad arms

To examine expression of INX-8(T239I D24N) mutant innexins encoded by *inx-8(rf)*, specific antibodies were used to label somatic (anti-INX-8) or the two classes of germline (anti-INX-21 or anti-INX-22) hemichannels. In wild-type hermaphrodites, clusters of gap junction puncta form between somatic DTC or sheath cells and each underlying germ cell (*Figure 2A*). In distal arms of wild type, loosely clustered sheath-germ cell gap junctions form more basally on germ cells; in *inx-8(rf)* distal arms, both INX-21 and INX-22 puncta tend to localize more laterally, where the sheath cells dip down between adjacent germ cells (*Figure 2B*). In the loop region (transition from meiotic pachytene to diplotene), large, higher-order clusters of gap junction puncta form in wild-type animals; these are absent in *inx-8(rf)* (*Figure 2C*). In wild-type proximal arms (diakinesis), developing oocytes exhibit extensive formation of INX-22 gap junction puncta, while INX-21 puncta become faint; in *inx-8(rf)* proximal arms, INX-22 and INX-8 are strongly but diffusely expressed, with little co-localization evident (*Figure 2D*). These results suggest there may be little gap junction coupling in *inx-8(rf)* proximal arms. We showed previously that sheath-oocyte gap junctions are endocytosed by the oocyte during ovulation (*Starich et al., 2014*). Embryos from *inx-8(0) inx-9(0)* rescued with *inx-8(rf)::gfp* show internalized GFP puncta, confirming that *inx-8(rf)* establishes at least some sheath-oocyte gap junctions in the proximal arm (*Figure 2–figure supplement 1*).

**Figure 2.**
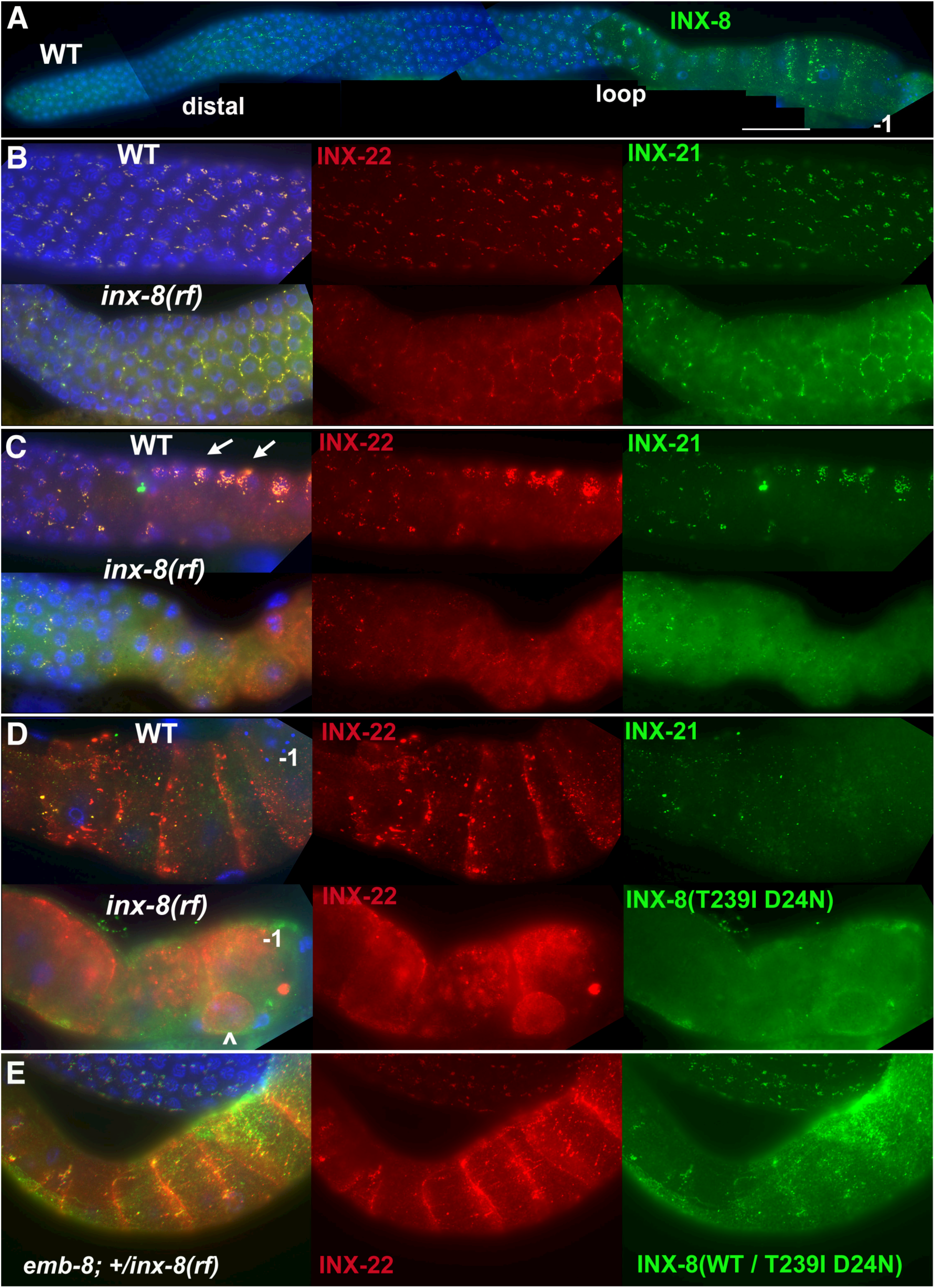
INX-8(T239I D24N) compromises gap junction formation between soma and germline. (A) Somatic INX-8 in wild-type (WT) gonads localizes to puncta representing soma-germline gap junctions where DTC or sheath cells overlie germ cells. (B) In WT distal gonad arms, gap junction puncta aggregate in small clusters associated with each germ cell; in *inx-8(rf)* distal arms, gap junction puncta localize more apically, where sheath dips down between germ cells, outlining the cell (honeycomb pattern). Localization of INX-21 and INX-22 define the two different classes of germline gap junction hemichannels, both of which include INX-14 subunits. (C) At the loop region, WT gap junction puncta form large higher-order aggregates (arrows) which are absent in *inx-8(rf)*. (D) In WT proximal arms, gap junction formation with INX-22-containing hemichannels is extensive; in *inx-8(rf)*, INX-22 and INX-8(T239I D24N) are strongly expressed but show little evidence of gap junction formation. “–1” indicates the most proximal oocyte; *inx-8(rf)* hermaphrodites frequently show undersized oocytes at this position as well (<). (E) Heterozygous *+/inx-8(rf)* shows little effect on gap junction localization in an *emb-8(hc69ts)* background. Antibodies specific to INX-8, INX-21, or INX-22 were used as indicated. DAPI-stained DNA in blue. Bar, 50 μm.

Reduced proximal arm expression of gap junctions is seen in *inx-8(0) inx-9(0); Ex[lag-2p::inx-8::gfp]* animals—simplified to *inx-8/9(0); Ex[inx-8(DTC+, Sh-)]*–which are rescued for germ cell proliferation by driving expression of INX-8::GFP in the DTC, but not sheath (Sh), using the *lag-2* promoter (*Starich et al., 2014*). These animals produce primarily dead embryos, many with Pod characteristics (*Table 1*). In contrast to *inx-8(rf)*, these embryos do not show evidence of *inx-8::gfp* endocytosis, consistent with absence of proximal arm sheath-germline gap junction coupling (*Figure 2–figure supplement 1*). *inx-8/9(0); Ex[inx-8(DTC+, Sh-)]* animals also show delayed onset of *gfp::lin41* expression, similar to *inx-8(rf)* (*Figure 1C; Figure 1–figure supplement table 2*).

We conclude that delayed gametogenesis in *inx-8(rf)* mutants arises from compromised soma-germline gap junction communication, due to altered channel properties that influence the formation and localization of gap junctions, especially in the loop and proximal arm. These regions are additionally implicated because DTC expression of wild-type INX-8 fails to rescue *inx-8(rf)* phenotypes (n>20).

### *inx-8(rf)* interacts with FAS mutants

Because *inx-8/9(0); Ex[inx-8(DTC+, Sh-)]* produces Pod-like embryos, and *inx-8(rf)* shows reduced proximal arm gap junction coupling similar to *inx-8/9(0); Ex[inx-8(DTC+, Sh-)]*, possible genetic interactions between *inx-8(rf)* and other Pod mutants were explored (*Figure 1–figure supplement table 1*). Conditional alleles of genes in the FAS pathway (*Figure 3A*) were used to construct double mutants with *inx-8(rf)*. Alleles included: *pod-2(ye60)*, a cold-sensitive (*cs*) allele of the *C. elegans* acetyl-CoA carboxylase ACC Type 1 (*Rappleye et al., 2003*); a temperature-sensitive (*ts*) allele of fatty acid synthase *fasn-1(g43) (Jaramillo-Lambert et al., 2015*); and *emb-8(hc69)*, a *ts* allele of an NADPH-cytochrome-P450 reductase, likely necessary for reduction of CYP-450 enzymes involved in fatty acid modifications (*Rappleye et al., 2003; Benenati et al., 2009*). Possible interaction between *pod-2(ye60cs)* and *inx-8(rf)* was difficult to assess–*ye60cs* grown at 15°C was quite sensitive to brief handling at room temperature. At a partially restrictive temperature (22°C), *fasn-1(g43ts); inx-8(rf)* and *emb-8(hc69ts); inx-8(rf)* double mutants exhibited markedly reduced brood sizes. Similar reductions in brood size were also apparent in *emb-8(hc69ts); +/inx-8(rf)*, suggesting *emb-8(hc69ts)* might be highly dose-sensitive to *inx-8(+)* levels (*Figure 1–figure supplement table 1*).

**Figure 3.**
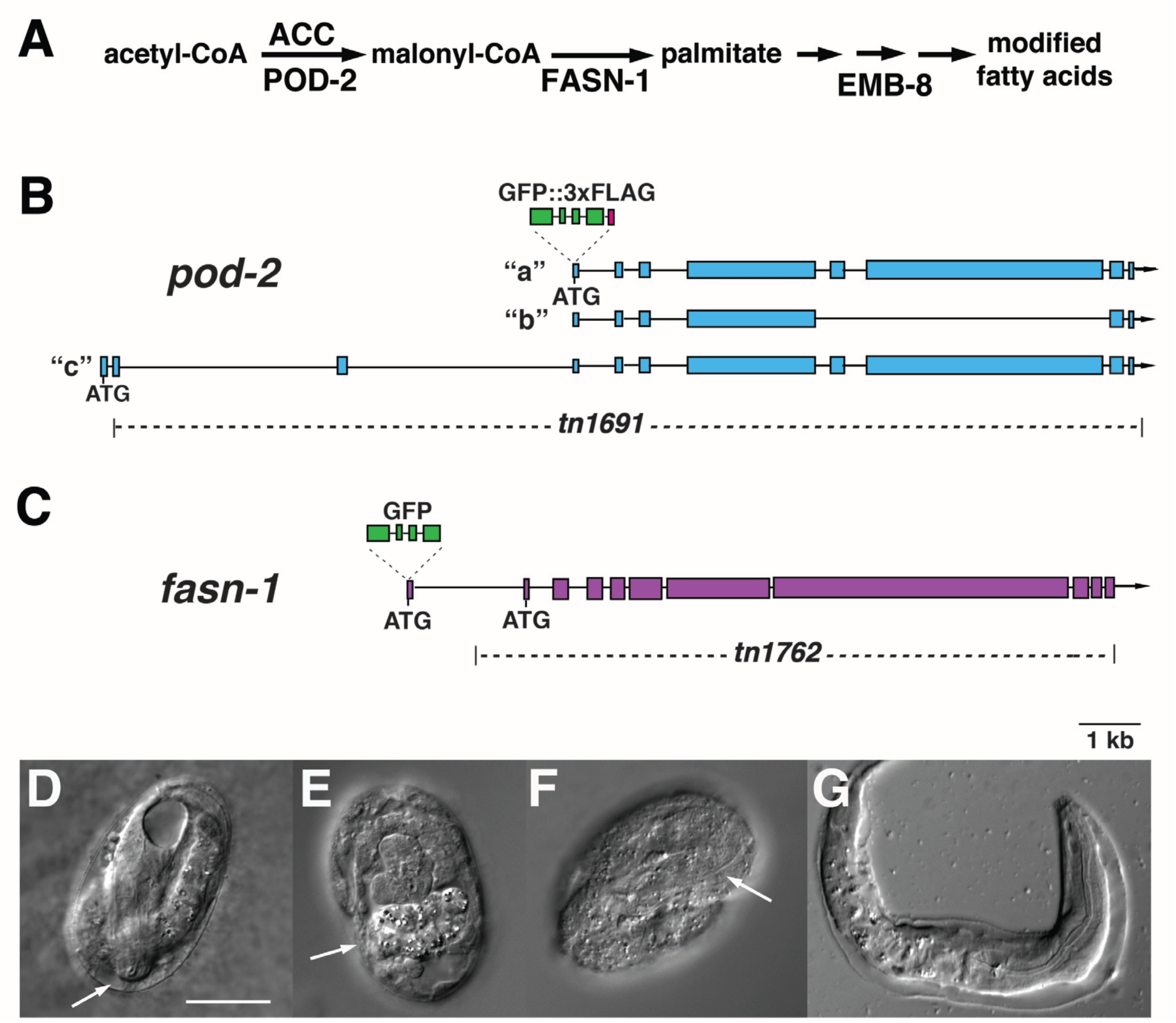
(A) Predicted assignment of FAS mutants in a simplified FAS pathway. Targeted deletions and GFP insertions are shown for *pod-2* (B) and *fasn-1* (C). Predicted isoforms for POD-2 are indicated. (D-G) Terminal phenotypes of (D) *pod-2(0)* and (E-G) *fasn-1(0)* mutant embryos from heterozygous *+/(0)* parents. Arrows in (D) and (F) indicate differentiated anterior pharynx; arrow in (E) indicates differentiated intestinal cells that formed gut granules. Bar, 20 μm.

### FAS pathway shifts from soma to germline after *pod-2*

Genetic interactions between *inx-8* and FAS genes could represent activity in the same or parallel pathways. If gap junctions are positioned in the FAS pathway, it might be possible to find a step within this pathway where gene function shifts from a somatic to a germline focus. *C. elegans* is particularly amenable to such an investigation because simple (highly repetitive) extrachromosomal arrays used to rescue gene function tend to express well in the soma, whereas arrays with more sequence complexity are generally required to prevent silencing of multi-copy gene expression in the germline (*Kelly et al., 1997*).

Previous characterization of *pod-2* implied rescue by a simple array (*Rappleye et al., 2003*); we likewise found that *pod-2(ye60cs)* is rescued by simple extrachromosomal arrays at 15°C (see Materials and methods). In contrast, 2 fosmids encoding *emb-8*, used individually to generate simple arrays, yielded 65 transformed lines, only one of which rescued *emb-8(hc69ts)* at 25°C; the array in this line was eventually silenced. For *fasn-1*, two fosmids used to generate 93 simple array lines and 20 complex array lines failed to rescue *fasn-1(g43ts)*. These results were consistent with a model in which FAS affecting embryonic development requires *pod-2* in the soma, but shifts to a germline requirement for *fasn-1* and *emb-8*.

To avoid misrepresentation of rescue results due to the special nature of a conditional allele, null alleles of *pod-2(tn1691)* and *fasn-1(tn1762)*–simplified to *(0)*–were generated using CRISPR-Cas9 methodology (*Figure 3B, C*) (Arribere *et al*., 2014). *pod-2(0*) embryos from *pod-2(0)/+* parents arrest late in embryogenesis (pretzel stage; *Figure 3D*); most *fasn-1(0)* embryos from *fasn-1(0)/+* parents shows signs of differentiation, and many arrest late in embryogenesis; occasional hatchlings emerge with probable hypodermal defects (*Figure 3E-G*). *pod-2(0)* was rescued by simple extrachromosomal arrays, supporting probable somatic function. Using a PCR-amplified genomic fragment we were able to rescue somatic but not germline function of *fasn-1(0)* with a simple or complex array (see Materials and methods). A genetically-balanced line with a complex array, *hT2/ fasn-1(0)*; *Ex[fasn-1(+); sur-5::gfp(+)]*, segregates healthy homozygous *fasn-1(0); Ex[fasn-1(+); sur-5::gfp(+)]* adults that lay almost exclusively Pod embryos (Table 1). (*sur-5::gfp* is a co-injection marker expressed in somatic cell nuclei (*Yochem et al., 19*98); corresponding extrachromosomal arrays can be lost mitotically to generate mosaics—see below). Gonad arms appear normal, and animals lay wild-type numbers of defective embryos (Table 1); many of these display polarity defects (4/14 symmetric first divisions) and exhibit extra nuclei in early embryos. Embryos typically arrest prior to morphogenesis (see below). We conclude this *fasn-1(+)* PCR fragment rescues somatic *fasn-1* function and is either silenced in the germline or lacks an element necessary for germline expression; in either case a germline requirement is indicated. In sum, rescue results are consistent with a fertility requirement for *pod-2* in the somatic gonad and *fasn-1* in the germline. This conclusion is supported by genetic mosaic and gene expression data described below. If *inx-8* gap junctions act in this pathway, it would position these junctions between *pod-2* and *fasn-1* as possible mediators of mal-CoA transfer from soma to germline.

### GFP::POD-2 and GFP::FASN-1 are tissue restricted

Endogenous, homozygous-viable, fertile GFP-labeled alleles of *pod-2* and *fasn-1* were generated to determine if patterns of localization are consistent with predicted foci of action. The GFP moiety in *gfp::pod-2(tn1765)* is expected to label all 3 predicted isoforms (*Figure 3B*) (*WormBase*). GFP::POD-2 expression is strongest in the intestine, and prominent in the hypodermis, gonadal sheath (especially proximal sheath), CAN neuron, and excretory duct (*Figure 4A-C*).

**Figure 4.**
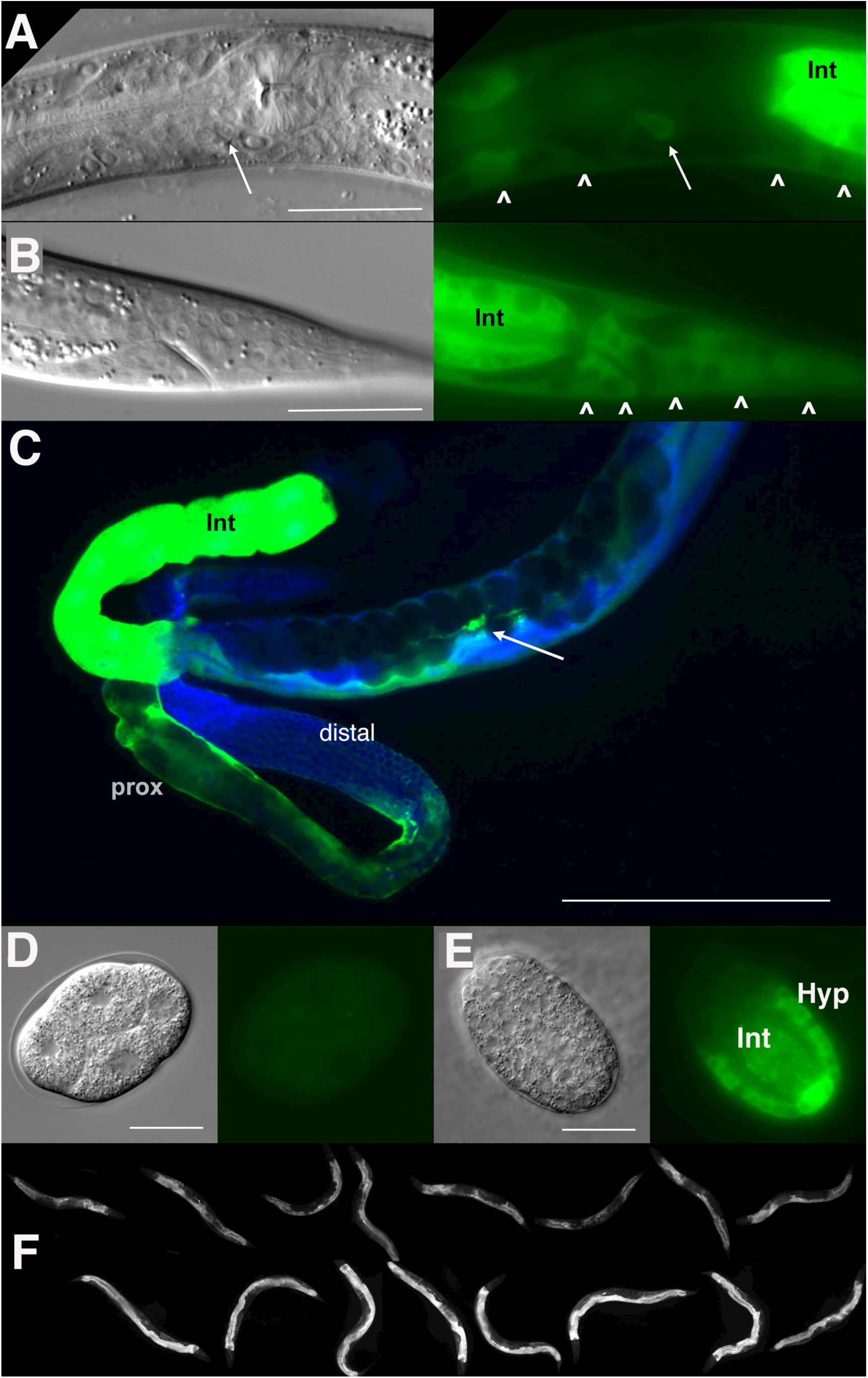
GFP::POD-2 expression. (A) Expression in the head includes hypodermis (carets) and excretory duct cell (arrow). Int, intestine. (B) Primary expression in the tail includes hypodermis (carets) and intestine (Int). Left panel DIC image, right panel GFP fluorescence in live animals. (C) Dissected animal stained with anti-GFP antibody shows abundant intestinal expression (Int) and strong expression in proximal sheath of exposed gonad arm (prox) that diminishes in the distal arm. No germline expression is seen in the distal arm where germ cells are revealed due to the absence of sheath coverage. Arrow indicates CAN neuron, one of the few neurons essential for viability in *C. elegans*. DAPI-staining of DNA in blue. (D, E) Earliest expression in embryos is evident in developing hypodermis and intestine. (A, B, D, E) DIC images in left panels, live GFP fluorescence in right panels. Bars, 20 μm, except (C) 200 μm. (F) GFP::POD-2 is responsive to starvation. Top row, animals briefly washed and transferred to bacteria-free NGM plates for 4 hours; lower row, animals similarly-treated but fed.

*gfp::fasn-1(tn1782)* labeled at the amino terminus (*Figure 3C*) is expressed in the hypodermis, gonad sheath (esp. proximal) and excretory duct; intestinal and CAN neuron expression are not evident (*Figure 5A-C*). This expression pattern partially overlaps with that of a *fasn-1* promoter::GFP reporter fusion, which was expressed in the intestine (*Lee et al., 2010*). WormBase predicts a single FASN-1 isoform, but a second in-frame ATG lies in Exon 2; intestinal FASN-1 could be derived from translational initiation at this site, which would not be revealed in our construct. Attempts to GFP-label FASN-1 downstream of this second ATG, or at the C-terminus, were unsuccessful. Neither *gfp::pod-2* nor *gfp::fasn-1* is expressed in the DTC.

**Figure 5.**
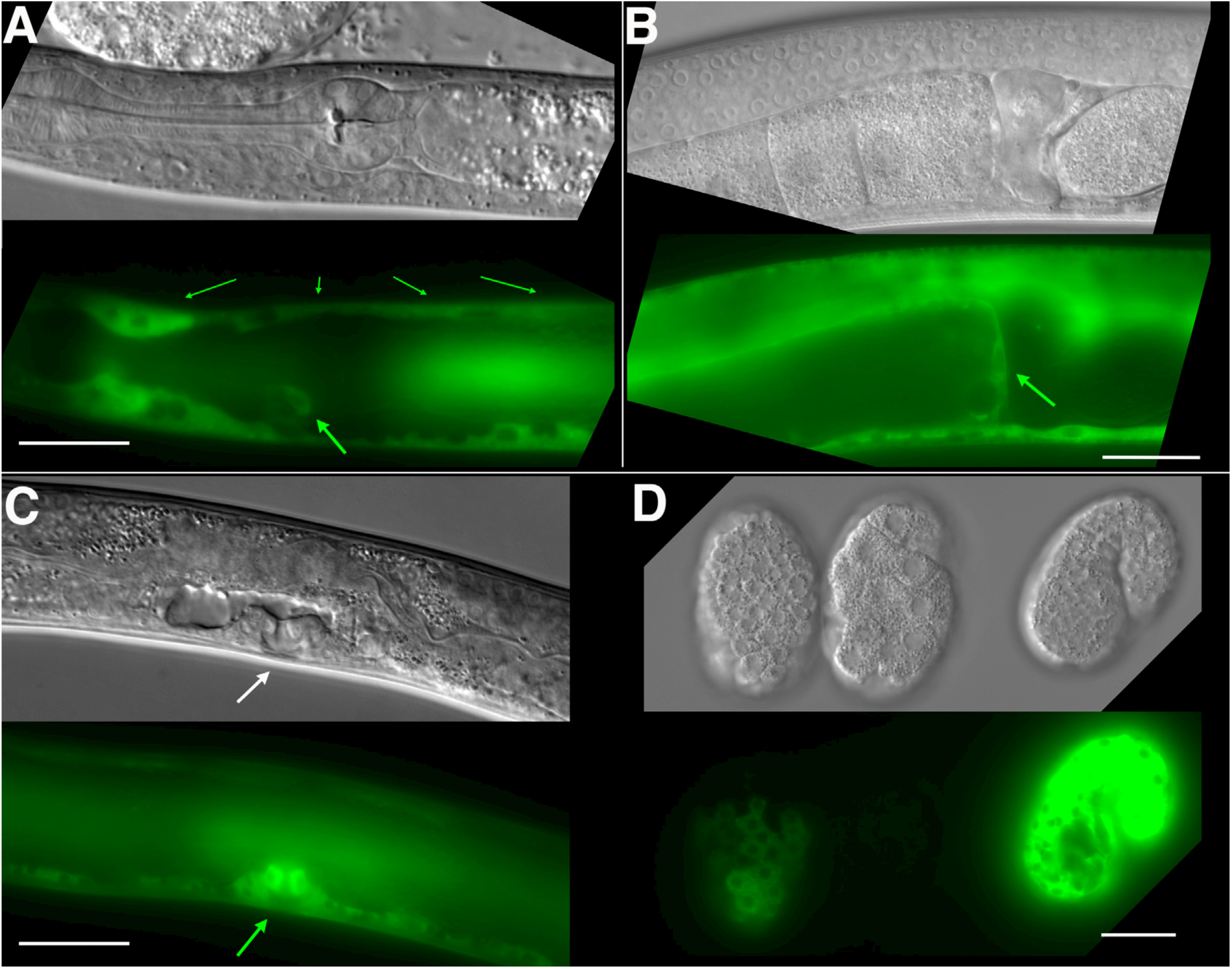
GFP::FASN-1 expression. (A) GFP::FASN-1 is expressed throughout the hypodermis (small arrows). Excretory duct cell expression is also indicated (larger arrow). (B) Arrow indicates proximal gonad sheath expression. (C) Arrow indicates developing vulva. (D) Earliest expression in developing embryos is seen in hypodermis. DIC images in upper panels, live GFP fluorescence in lower panels. Bars, 20 μm.

Germline expression for either GFP-tagged gene was not detected (*Figure 4C; Figure 5B*). Anti-GFP staining of dissected gonads failed to detect germline GFP::POD-2 expression (*Figure 4C*). The earliest detectable GFP::FASN-1 in embryos appears in developing hypodermis (*Figure 5D*); GFP::POD-2 is prominent in developing hypodermis and intestine (*Figure 4D, E*). GFP::POD-2 expression decreases in response to starvation (*Figure 4F*).

Expression of both genes is consistent with primarily somatic functions. Germline *fasn-1* activity may derive from low levels of maternal expression, though its presence is confirmed by the late arrest phenotype of *fasn-1(0)* embryos derived from *+/fasn-1(0)* parents (*Figure 3E-G*).

### INX-8 channels influence effects of overexpressed malonyl-CoA on downstream FAS mutations

If *inx-8* is required for transit of mal-CoA from soma to germline through gap junctions, the fertility of mutants downstream of *pod-2* in the FAS pathway could be compromised by inclusion of INX-8(T239I D24N) into somatic hemichannels. Expression of an *Ex[pod-2(+); sur-5::gfp]* simple array increases the brood size of *emb-8(hc69ts)* at 22°C. This increase is dependent on INX-8 hemichannel integrity—*inx-8(+)* homozygotes facilitate the increase more effectively than *+/inx-8(rf)* heterozygotes (*Figure 6A*). Importantly, anti-INX-8 and anti-INX-22 co-localization in *emb-8(hc69ts); +/inx-8(rf)* dissected gonads is indistinguishable from wild type, implying that cell adhesion is not affected (*Figure 2E*).

**Figure 6.**
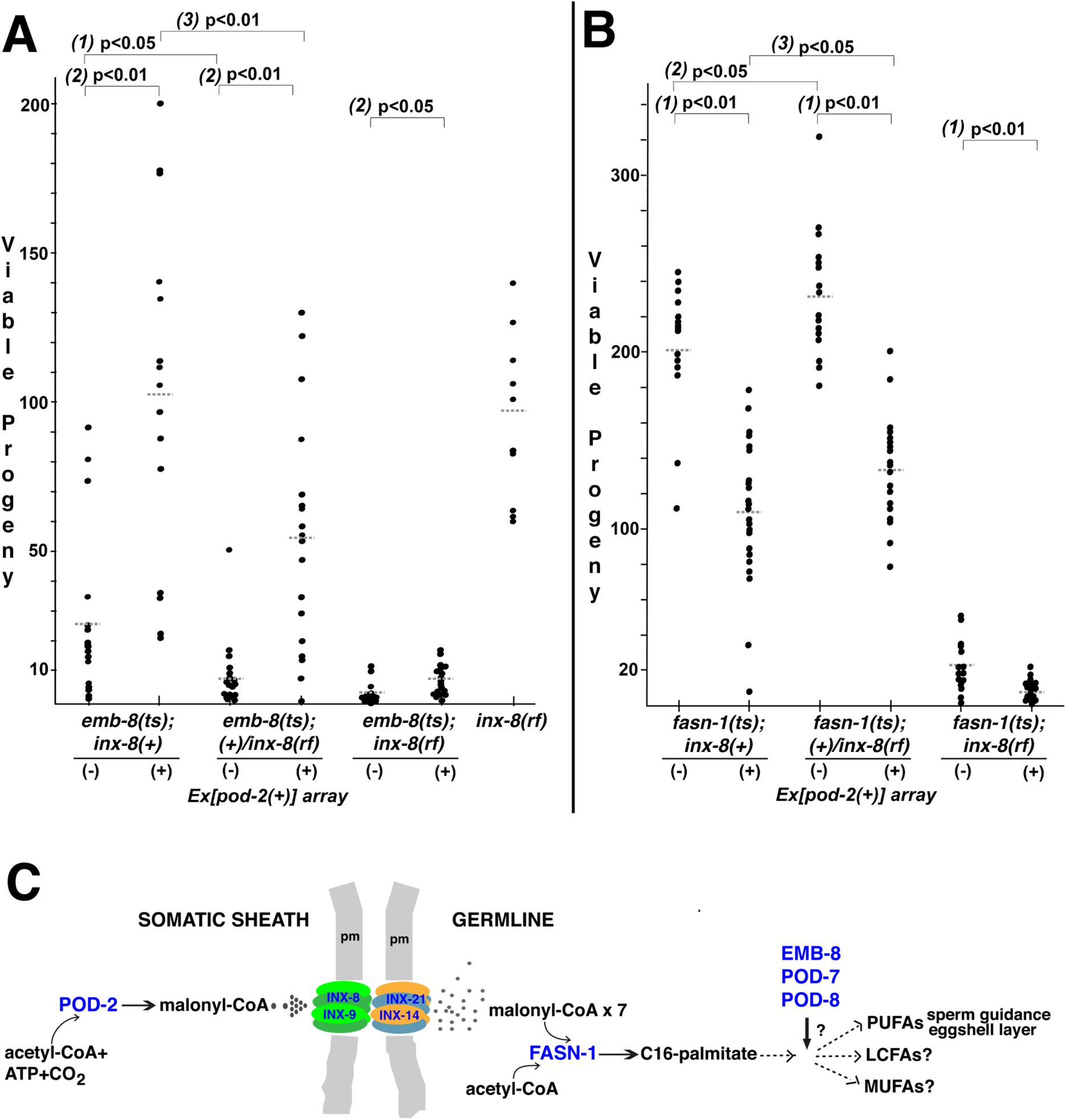
Wild-type INX-8 facilitates effects of *Ex[pod-2(+)]* overexpression on conditional mutations downstream in the fatty acid synthesis (FAS) pathway. Brood size was measured as the number of viable progeny produced. The balancer *mIs11* provided *inx-8(+)* for all strains listed. (A) Embryonic lethality in *emb-8(ts)* mutants at 22°C is more severe in heterozygous *+/inx-8(rf)* than *inx-8(+)* backgrounds *(1)*. Expression of *Ex[pod-2(+)]* in *emb-8(ts)* significantly rescues embryonic lethality *(2)*. Rescue of *emb-8(ts)* by *Ex[pod-2(+)]* is also dependent on *inx-8(+)* copy number *(3)*. (B) Expression of *Ex[pod-2(+)]* in *fasn-1(ts)* mutants at 20°C reduces brood size *(1). fasn-1(ts); inx-8(+)* homozygotes show greater embryonic lethality than *fasn-1(ts); (+)/inx-8(rf*) heterozygotes in the absence *(2)* or presence *(3)* of *Ex[pod-2(+)]*. Averages indicated with dotted lines. Probability was calculated using Student’s 2-tailed *t*-test. (C) Enzymes (purple) implicated in germline FAS. Malonyl-CoA transits from soma sheath to germline through gap junction channels comprised of INX-8/9 (soma) and INX-14/INX-21 (germline) hemichannels. Germline FASN-1 uses malonyl-CoA to synthesize palmitate, which may be further extended and modified to produce other long-chain fatty acids. EMB-8 (NADPH-dependent CYP450 reductase) and POD-7/8 (CYP450s) are necessary to synthesize the lipid-rich (inner) layer of the eggshell and support polarity establishment (*Rappleye et al., 2003; Benenati et al., 2009*); the composition of lipid layer fatty acids is unknown. *de novo* PUFA synthesis is required in the germline for embryonic viability, and is implicated in sperm guidance (*Kubagawa et al., 2006; Watts et al., 201*8). The identities of other long-chain fatty acids that may be synthesized in the germline are unknown. PUFAs, polyunsaturated fatty acids; LFAs, long-chain fatty acids; MUFA, monounsaturated fatty acids; pm, plasma membrane. **Source data 1.** Viable progeny counts for the Figure 6 panels A, B.

In contrast to *emb-8(hc69ts)*, expression of *Ex[pod-2(+); sur-5::gfp]* in *fasn-1(g43ts)* significantly <u>reduced</u> brood size (p<0.01), and brood size reduction is more severe in homozygous *inx-8(+)* than heterozygous *+/inx-8(rf)* backgrounds (ρ<0.05) (*Figure 6B*). This result likely relates to the mechanism of FASN production of fatty acids—acetyl-CoA (FA chain initiation) and mal-CoA (chain elongation) load at a common, non-selective acyl-CoA binding site and are regarded as competitive inhibitors (*Cox and Hammes, 1983; Chang and Hammes, 1990*). Increasing the ratio of mal-CoA to acetyl-CoA may inhibit FASN-1 enzyme by interfering with chain initiation. Both wild-type and enhanced levels of mal-CoA delivered through wild-type INX-8 hemichannels appear to inhibit *fasn-1(g43ts)*; this mutant form of FASN may be especially sensitive to inhibition by higher levels of mal-CoA. Importantly, heterozygous *+/inx-8(rf)* channels ameliorate this inhibition, likely by reducing mal-CoA transfer.

Enhancement by *inx-8(+)* hemichannels of both a positive *(emb-8)* and a negative *(fasn-1)* effect on FAS mutations downstream of overexpressed *pod-2(+)* strongly argues that *inx-8* acts in the same pathway as *pod-2*, and is consistent with gap junctions acting as conveyors of mal-CoA from soma to germline (*Figure 6C*).

### Genetic mosaic analyses associate *pod-2* sheath cell loss with the Pod phenotype

To verify a somatic *pod-2* focus of action, genetic mosaic animals derived from *pod-2(0); Ex[pod-2(+); sur-5::gfp]* were sought. 17/833 adult hermaphrodites were identified that produced dead eggs but no viable progeny; these adults were examined for cell lineage loss of *Ex[pod-2(+); sur-5::gfp]*. Animals fell into two classes: (1) eight animals lost *Ex[pod-2(+); sur-5::gfp]* from most or all gonad sheath cells, and (2) nine animals lost *Ex[pod-2(+); sur-5::gfp]* from the intestine (*Figure 7A*).

**Figure 7.**
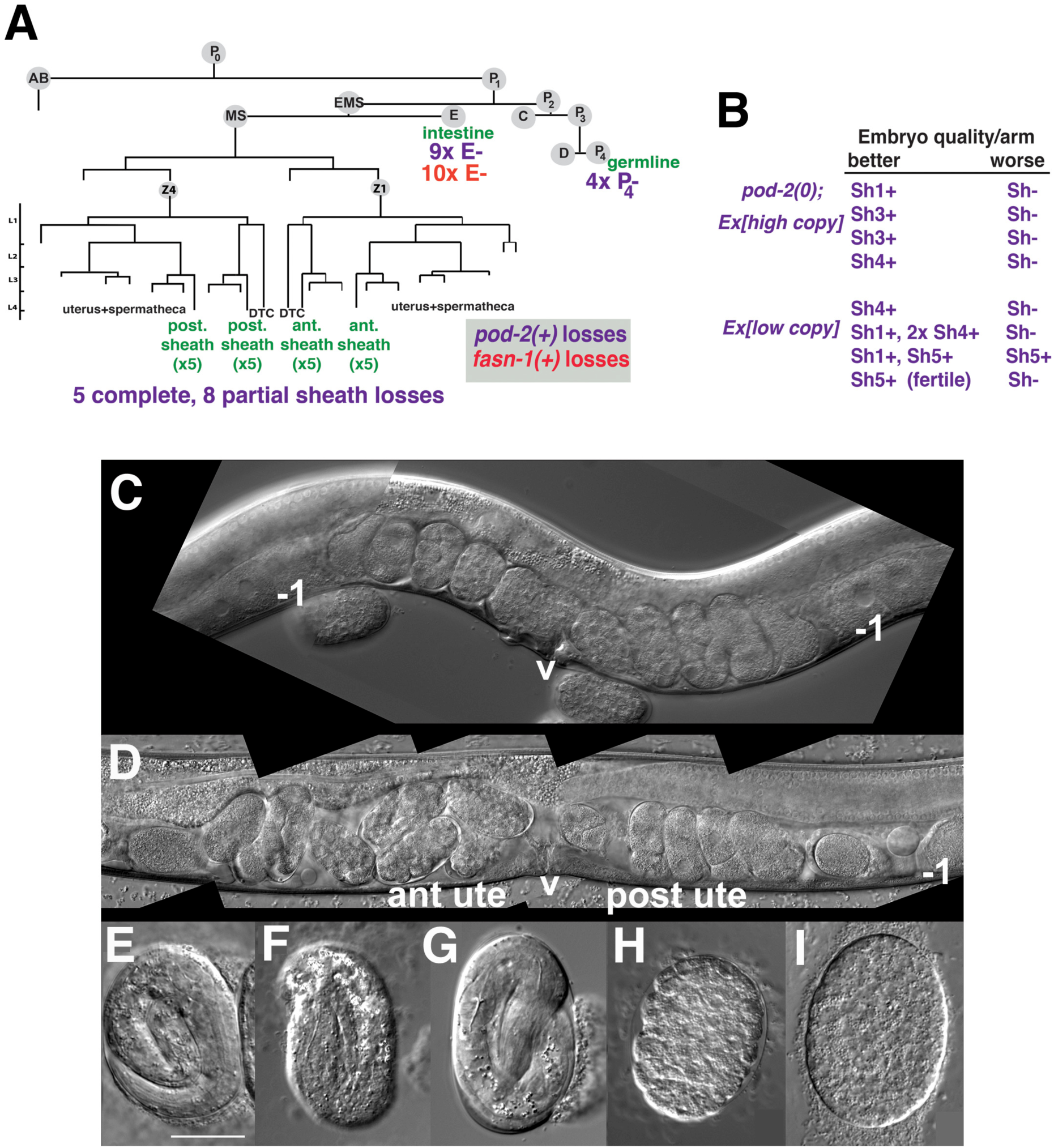
Genetic mosaic analyses of *pod-2* reveal a mal-CoA requirement in somatic gonad sheath. (A) Abbreviated cell lineage of *C. elegans*. Mosaics characterized correspond to loss of *sur-5::gfp* expression in the germline (P_4_), intestine (E), or somatic gonad. Five *pod-2(0); Ex[pod-2(+); sur-5::gfp]* animals lost the array from all somatic gonad sheath cells and produced no viable embryos. (B) Eight animals showed differences in array retention in each gonad arm with phenotypic consequences. Arrays were generated by microinjecting cosmid W09B6 *[pod-2(+)]* at 4 ng or 20 ng/μl (low/high copy). (C) Uterus of non-mosaic *pod-2(0); Ex[pod-2(+)—*high copy] hermaphrodite; developmental progression of embryos from each gonad arm is comparable. (D) Mosaic *pod-2(0); Ex[pod-2(+)—*high copy] with array lost from posterior gonad arm sheath but retained in a single Sh4 cell of anterior arm. Anterior arm embryos have more distinct eggshells and develop to later stages. (E-H) Embryonic development depends on somatic levels of mal-CoA. Shown are terminal-stage *pod-2(0)* embryos derived from parents of designated genotypes. (E) Pretzel-stage embryo from *+/pod-2(0)*. (F) 3-fold stage embryo from *pod-2(0); Ex[pod-2(+)— low copy]*, with array maintained in soma but lost in the germline (P4–). (G) Pretzel-stage embryo from P4(–) mosaic *pod-2(0); Ex[pod-2(+)—high copy]*. (H) Embryo from mosaic *pod-2(0); Ex[pod-2(+)—low copy]* with uncharacterized somatic loss of array, but array retained in the germline (P4+). This parent produced exclusively dead embryos. (I) Typical terminal undifferentiated *fasn-1(0)* Pod embryo from *fasn-1(0); Ex[fasn-1(+)]*. ant ute, anterior uterus; post ute, posterior uterus; v, vulva; –1, most proximal oocyte. Bar, 20 μm.

Within the first class of mosaics, five animals lost the array in all sheath cells in both gonad arms. Three other animals exhibited a difference in phenotypic severity between gonad arms—embryos derived from one arm developed to later multicellular stages *in utero* (*Figure 7B-D*). Differences were coincident with loss of the array from all sheath cells in the more severely affected arm, versus retention of the array in a single sheath cell of the better arm. In subsequent screening, five more mosaics were identified that displayed phenotypic disparity between the two gonad arms; in one example, one arm produced defective embryos while the better arm gave rise to wild-type embryos (producing ≥40 viable progeny). These results indicate that the focus of *pod-2(0)* rescue is autonomous to each gonad arm, consistent with the interpretation that *pod-2* expression in the somatic sheath is essential to embryonic viability.

In subsequent screening for germline (P_4_ –) mosaics, four animals were identified that maintained the array in the soma but lost the array in all embryos (*Figure 7A*). The terminal stages of embryonic development resembled those of *pod-2(0)* embryos derived from *+/pod-2(0)* parents, i.e. embryos developed to 3-fold or pretzel stages but failed to hatch (*Figure 7E-G*). These stages are more advanced than terminal stages of many dead embryos derived from parents with somatic (but not germline) loss of the rescuing array; such embryos fail to differentiate (*Figure 7H*). This early developmental arrest resembles terminal-stage Pod embryos from *fasn-1(0); Ex[fasn-1(+))*] (compare panels H and I in *Figure 7*). We conclude that somatic *pod-2(+)* supplies sufficient levels of mal-CoA to the germline not only to establish a wild-type 1-cell embryo, but also to support continued development, presumably until zygotic *pod-2* expression becomes adequate for completion of embryogenesis. These studies establish that metabolites essential for embryonic development can be produced in the somatic gonad and provided to the oocyte through gap junctions.

### Intestinal loss of *pod-2*, but not *fasn-1*, results in a germline starvation response

The other class of *pod-2(0)* genetic mosaics producing dead eggs in our original screen lost *Ex[pod-2(+); sur-5::gfp]* from the intestine (E–) (*Figure 7A*). E(–) mosaic animals were smaller than wild type, and gonad arms were severely reduced in size (*Figure 8A*). Only the most proximal oocytes showed appreciable growth. E(–) mosaic gonad arms resemble those seen in gonads undergoing the oocyte starvation response, or adult reproductive diapause (ARD), a process induced by starvation during the L4 larval stage proposed to maintain germline stem cells for repopulation of the gonad upon the resumption of feeding (*Angelo and Van Gilst, 2009; Seidel and Kimble, 2011*). During ARD, germline stem cells cease dividing, the gonad shrinks due to extensive apoptosis of differentiated germ cells, and growth is largely restricted to the most proximal oocyte.

**Figure 8.**
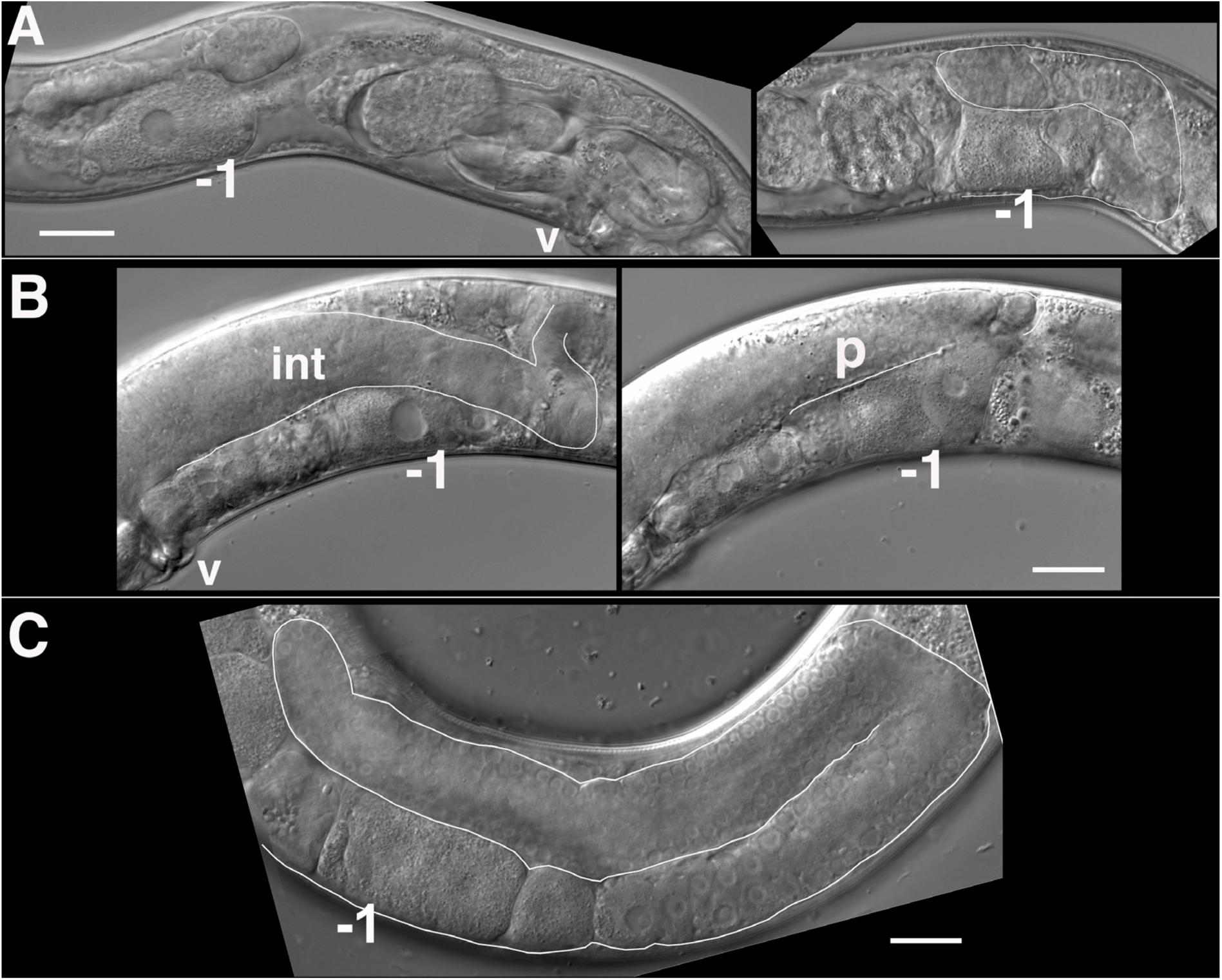
Mosaic loss from the intestine of *pod-2*, but not *fasn-1*, leads to an ARD-like gonad arm. (A) E(–) genetic mosaic *pod-2(0); Ex[pod-2(+); sur-5::gfp]* gonad arms (from same animal). Gonad arm on right partially outlined. (B) Two focal planes of a presumptive P_1_(–) *pod-2(0); Ex[pod-2(+); sur-5::gfp]* mosaic with constipated intestine. Upper plane (left) with partially outlined intestine; lower plane (right) shows underlying distal gonad arm. int, intestine; p, pachytene region; v, vulva. (C) E(–) genetic mosaic *fasn-1(0); Ex[fasn-1(+); sur-5::gfp]* gonad arm. –1, most proximal oocyte. Bars, 20 μm.

The *C. elegans* intestine is rich with lipid droplets and the source of yolk, and regarded as a major site of FAS (*Watts and Ristow, 2017*). A genetic pathway in *C. elegans* required for intestinal apical polarity establishment leading to proper lumen formation has been described; genes in this pathway, including *pod-2*, encode enzymes required for glycosphingolipid synthesis (*Zhang et al., 2012*). Lipid biosynthesis is required throughout larval growth, coincident with membrane expansion. The identification of adult mosaics lacking *pod-2(+)* in the intestine was therefore surprising. We subsequently identified two presumptive P_1_(–) *pod-2* mosaics, predicted to have lost *pod-2(+)* function in the intestine, somatic gonad, germline, many muscle and hypodermal cells, among others (*Figure 7A*). The most striking phenotype distinguishing these animals from E(–) mosaics was an extremely constipated intestine (*Figure 8B*). The absence of this constipation in E(–) mosaic animals implies that loss of *pod-2(+)* from the intestine does not prevent the processing of bacteria through the intestinal lumen. (Both P_1_ mosaics showed tissue extruded from the rectum, which may be the cause of constipation.)

Although *gfp::fasn-1(tn1782)* is not detected in the intestine, we asked if there is a functional role for *fasn-1* similar to *pod-2*. As described above, homozygous *fasn-1(0); Ex[fasn-1(+); sur-5::gfp]* progeny from heterozygous parents produce Pod embryos but exhibit no other phenotype; within this class of progeny we screened for E(–) mosaics. Ten (of 800) such animals were identified (*Figure 7A*); in stark contrast to *pod-2*, their gonad arms appeared normal. These animals displayed no overt defects outside of Pod embryo production (*Figure 8C*). Because maternal FASN-1 rescues *fasn-1(0)* homozygotes to later embryonic stages, an early intestinal function for *fasn-1* equivalent to *pod-2* could be rescued maternally. However, the surprising conclusion from this mosaic analysis is that there is no significant intestinal requirement for zygotic *fasn-1* in larval and adult stages. The implication is that the primary function of mal-CoA produced in the intestine is not as substrate for intestinal FASN-1.

### Is malonyl-CoA required in other germline processes?

*inx-8(rf)* and *inx-8/9(0); Ex[inx-8(DTC+, Sh-)]* share delayed oogenesis and a reduction in gap junction coupling, germ cell proliferation, and brood size. *inx-21(0)* mutants are sterile, and *inx-22(0)* mutants are fertile (*Starich et al., 2014*); therefore mal-CoA required for embryos must transit through gap junction channels comprised of INX-14/INX-21 hemichannels, which are expressed throughout the distal arm (*Figure 2B*). We asked if mal-CoA levels might influence phenotypes preceding embryogenesis, such as oocyte production.

Supplying mal-CoA transiently by microinjection (0.5-10 mM) into *inx-8/9(0); Ex[inx-8(DTC+, Sh-)]* L4 gonad arms failed to ameliorate phenotypes such as Pod-embryo production or low brood size. Ectopic expression of GFP::POD-2 in the germline in *inx-8/9(0); Ex[inx-8(DTC+, Sh-)]* or *inx-8(rf)* genetic backgrounds also failed to improve phenotypes, including delayed gametogenesis (*Figure 7–figure supplement 1*). However, whether endogenous mal-CoA levels are altered under these conditions is uncertain.

Effects of global mal-CoA levels were indirectly examined by comparing higher or lower *pod-2(+)* copy number rescue of *pod-2(0)*; numbers of dead eggs and larvae were summed to reflect total embryo output (Table 1). Lower copy number rescue coincided with reduced embryo output (84% of wild-type) and lower proportion of viable larvae within a brood (∼37% of total embryos) (Table 1). Embryo output may therefore by impacted by *pod-2* expression; however, somatic cells outside the gonad could be responsible. We therefore asked specifically if FAS is required within the germline. Somatically-rescued *fasn-1(0); Ex[fasn-1(+); sur-5::gfp]* embryo output is comparable to wild-type (94%); however, embryo output in the double mutant *fasn-1(0); inx-8(rf); Ex[fasn-1(+); sur-5::gfp]* is only ∼50% of *inx-8(rf)* alone (Table 1). The FAS pathway may therefore play a germline role affecting embryo output. Surprisingly, *inx-8(rf)* partially suppresses the embryonic lethality of *fasn-1(0); Ex[fasn-1(+); sur-5::gfp]* (Table 1). Possibly the smaller gonad arms of *inx-8(rf)* may concentrate levels of FASN-1 or mal-CoA. Alternatively, *Ex[fasn-1(+); sur-5::gfp]* germline expression may be de-silenced in this genetic background. Further study is needed to address the nature of this suppression.

## Discussion

The genetic analysis of mutant *inx-8* soma-germline gap junction channels has led us to three conclusions that have significance beyond the *C. elegans* model system: (1) Mal-CoA produced in the somatic sheath is delivered through gap junctions to the germline; this mal-CoA is necessary to establish a proper one-cell embryo and to support continued embryogenesis. To our knowledge, this segregation of the cellular source of mal-CoA from the eventual cell type in which it is used as substrate is novel. As the rate-limiting step in FAS, mal-CoA production is an attractive control point to tie aspects of germ cell developmental progression to somatic control. (2) *pod-2* expression (and presumably mal-CoA production) is exceptionally robust in the intestine and essential for normal germline development. In the absence of intestinal POD-2, the germline exhibits phenotypic hallmarks of germline ARD despite the presence of a replete food source. By contrast, intestinal *fasn-1* expression is not similarly required. (A possible caveat is that maternal *fasn-1* in the intestine may be sufficient to reach adulthood, which would be surprising.) In light of the abundant expression level of *gfp::pod-2* in the intestine, and our finding that mal-CoA can be directionally transferred intercellularly, the question arises as to whether the intestine provides mal-CoA to other cells. Alternatively, a deficit of mal-CoA production in the intestine might initiate a systemic starvation response. Recently, *Gerisch et al.*, (2020) have shown that the HLH-30/TEFB transcription factor functions as a master regulator of the ARD state. *hlh-30* mutants fail to effect the system-wide changes promoting long-term survival and recovery associated with ARD. ARD longevity does not appear to rely on fatty acid metabolism. However, entry of *hlh-30* mutants into the ARD state appears normal; therefore effects of decreased mal-CoA expression in the intestine may lie upstream of *hlh-30* activity. (3) A pronounced delay in gametogenesis along with reduction in brood size results from reduced soma-germline gap junction communication; it is unknown if this delay derives from generally slower mitotic or meiotic progression, or from a developmental block at a particular meiotic stage. The nature of potential molecules passing through gap junctions that are responsible for timely meiotic progression is unknown, but mal-CoA has thus far not been implicated.

Our results are consistent with other findings in *C. elegans* related to fatty acid metabolism in oocytes. *pod-2*/ACC requires post-translational covalent attachment of biotin for enzymatic activity, and *bpl-1* (*b*iotin *p*rotein *l*igase-1) is the *C. elegans* ortholog of the necessary holocarboxylase synthetase (*Watts et al., 2018*). Mal-CoA provided in the bacterial diet enables *bpl-1* mutants to develop to adulthood, but *de novo* synthesis of polyunsaturated fatty acids (PUFAs) is required for embryonic viability (*Watts et al., 2018*). Our results indicate that mal-CoA supplied for PUFA synthesis (and other FAS) in the early embryo is derived from the somatic sheath via gap junctions. PUFA-dependent prostaglandin signaling from oocytes influences sperm guidance, and reduction of *inx-14* function disrupts this signaling (*Kubagawa et al., 2006; Edmonds et al., 2011*). We suggest that reduced levels of mal-CoA delivered to the oocyte through mutant INX-14*-*containing gap junctions may reduce prostaglandin synthesis and its signaling function for sperm migration.

At least one route outside of gap junctions may deliver mal-CoA to the germline early in its development. The primordial germ cells Z2 and Z3 associate with a pair of intestinal cells during embryogenesis; contacts are lost before hatching (*Sulston et al., 1983; Abdu et al., 2016*). Therefore an early germline requirement for mal-CoA could be fulfilled directly from the intestine. Loss of such a function would be encompassed within the ARD-like phenotype seen in *pod-2* E(–) mosaics.

In addition to its metabolic function in FAS, mal-CoA has established roles as a regulatory molecule. It inhibits beta-oxidation of fatty acids by mitochondria. Inhibitory mal-CoA derives from a second acetyl-CoA carboxylase (ACC2), associated with the mitochondrial membrane. Manipulation of mal-CoA levels in the mouse hypothalamus regulates food intake (Wolfgang and Lane, 2006). Regulation is effected through the brain-specific CPT1-c, via a mechanism distinct from the enzymatic activity of other CPT1 enzymes (*Sierra et al., 2008; Reilly and Mak, 2012*). Little information regarding the *C. elegans* ACC2 is known; somatically-delivered mal-CoA could potentially influence germline mitochondrial function.

Malonylation as a post-translational lysine modification has recently been described (*Peng et al., 2011*). Malonylation may inhibit target proteins, such as the glyocolytic enzymes glyceraldehyde 3-phosphate dehydrogenase and pyruvate kinase (*Kulkarni et al., 2017*). Malonylation of mTOR reduces mTORC1 complex activity, decreasing endothelial cell proliferation and reducing angiogenesis in a human umbilical model (*Bruning et al., 2018*). Together these studies and others suggest a potentially pervasive role for mal-CoA in sensing and responding to nutritional status. One trait associated with rapid tumor growth is an increase in FAS, and FASN has been explored as a cancer therapy target (*Menendez et al., 2007*). Possibly the ability of mal-CoA to transit through gap junctions might sustain FAS during tumor progression. The discovery that mal-CoA transits through gap junctions from soma to germline may be indicative of other unappreciated intercellular roles for this important molecule.

## Materials and methods

### Strains

*C. elegans* strains were grown on standard NGM media (plus Nystatin, 6.25 mg/l) with *E. coli* strain OP50 as food source. Temperatures used for conditional mutants are indicated in main text. In addition to the wild-type strain N2, the following alleles were used: Chr. *I*: *fasn-1(g43ts); fasn-1(tn1762null); fasn-1(tn1782[gfp::fasn-1]); lin-41(tn1541[gfp::tev::s-tag::lin-41]);* Chr. *II*: *pod-2(ye60cs); pod-2(tn1691null); pod-2(tn1765[gfp::tev::3xFLAG::pod-2]);* Chr. *III*: *emb-8(hc69ts); dpy-17(e1295);* Chr. *IV*: *inx-8(tn1474null) inx-9(ok1502null); inx-8(tn1513) inx-9(ok1502); inx-8(tn1513 tn1555) inx-9(ok1502)—*a.k.a. *inx-8(rf)* in main text. Balancers used included: *tmC18 [dpy-5(tmIs1236)] I; hT2[bli-4(e937) let-?(q782) qIs48] (I:III); mIs11 IV.* Extrachromosomal arrays used included: *tnEx195[inx-8(+) inx-9(+); sur-5::gfp]; tnEx205[lag-2p::inx-8::gfp; str-1::gfp]; tnEx212[W09B6-pod-2(+)—20 ng/μl—hi copy; sur-5::gfp]; tnEx218[fasn-1(+); sur-5::gfp]; tnEx219[W09B6-pod-2(+)—4 ng/μl—lo copy; sur-5::gfp];* and *tnEx221[inx-8(tn1513 tn1555); str-1::gfp].*

### Microinjection rescue of FAS mutants with extrachromosomal arrays

The cosmid W09B6 was used at 50 ng/μl (2 lines) or 20 ng/μl (1 line) with *sur-5::gfp* (75 ng/μl) to rescue *pod-2(ye60cs). sur-5::gfp* expression was faint for both lines obtained with 50 ng/μl and these were not further characterized. *pod-2(tn1691null)* was rescued by microinjection using W09B6 at 4 ng/μl (5 lines) or by crossing in *tnEx212*. Fosmids WRM0613bD07(10 ng/μl, or 50 ng/μl—1 line that silenced) and WRM067cB03 (25 ng/ul) were used to attempt rescue of *emb-8(hc69ts)*. Numerous conditions using fosmids WRM0612bD08 and WRM0614aH05 in simple or complex arrays were used to try to rescue *fasn-1(g43ts)* without success. However, FASN enzyme functions as a dimer; formation of dimers between mutant (*g43*) and wild-type monomers might inhibit phenotypic rescue. After *fasn-1(tn1762null)* was generated, somatic but not germline rescue was obtained using a *fasn-1* genomic PCR product forming simple arrays (1 ng/μl + *sur-5::gfp-75* ng/μl, at least 2/6 lines segregated *fasn-1(0); Ex[fasn-1(+); sur-5::gfp(+)]* animals that laid only Pod embryos), or a complex array (1 ng/μl + 20 ng/μl *sur-5::gfp* + 100 ng/μl salmon sperm,1 line from 30 injected animals) as described in the main text.

### Recombinant DNA constructs for isolation of new *pod-2* and *fasn-1* alleles

Standard CRISPR/Cas-9 methods were used to generate deletions and GFP insertions in *pod-2* and *fasn-1.* For deletions, 100-nt repair templates were synthesized (Ultramers; Integrated DNA Technologies, Coralville, IA, USA) and used in a *dpy-10(cn64)* co-conversion strategy previously described (*Arribere et al., 2014*). Injection mixes included: pDD162 (Cas9), 50 ng/μl; 25 ng/μl (*pod-2*) or 10 ng/μl (*fasn-1)* each single guide; Ultramer repair template, 500 nM; *dpy-10* single guide (pJA58), 25 ng/μl; *dpy-10(cn64)* Ultramer A2-2F-827, 500nM. Single guides were made by annealing oligo pairs and cloning into pRB1017 *(Arribere et al., 2014*): Deletions were verified by PCR. The PCR fragment used to rescue *fasn-1(tn1762null)* was amplified from genomic DNA using primers 5’-tacgtcaaacatccgtttgtcaacgtcac-3’ and 5’-ttcttctcctcgtcctttgatacgaagatc-3’.

Generation of GFP insertions into *pod-2* using pDD282 followed established protocols (*Dickinson et al., 2015*). Injection mix included the single guide (10 ng/μl), repair template (10 ng/μl), pDD162 (50 ng/μl), and *sur-5::gfp* (80 ng/μl). The insertion was verified by sequencing PCR products generated with primers positioned outside the homology arms.

For *gfp::fasn-1*, we were unable to isolate 3’ insertions using the Dickinson method (*Dickinson et al., 2015*). This method relies on isolation of candidates based on the Roller phenotype, and because *fasn-1* has been implicated in molting, we turned to a co-conversion approach (*Arribere et al., 2014*). A repair template was generated that included the GFP sequence from pDD282, and *fasn-1* homology arms on either side. The injection mix contained pDD162, 50 ng/μl; *fasn-1* single guide 10 ng/μl; *gfp::fasn-1* repair template, 20 ng/μl; *dpy-10* single guide (pJA58), 25 ng/μl; *dpy-10(cn64)* Ultramer A2-2F-827, 500nM^;^ and pCFJ190, 2.5 ng/μl. Rol candidates were screened directly for GFP expression. The insertion was verified by sequencing PCR fragments generated using primers placed outside of the repair template sequence.

*mex-5p::gfp::pod-2::tbb-2-3’UTR* was constructed by Gibson assembly of component PCR products representing the *mex-5* promoter, the “a” isoform of *gfp::pod-2*, and the tbb-2-3’UTR to prevent germline RNA degradation.

### Immunofluorescence

Antibody staining of dissected gonads was as previously described (*Starich et al., 2014*). DIC and fluorescent images were acquired on a Zeiss (Thornwood, NY, USA) motorized Axioplan 2 microscope with either a 40x Plan-Neofluar (numerical aperture 1.3), a 63x Plan-Apochromatic (numerical aperture 1.4), or 100x PlanApochromatic (N.A. 1.4) objective lens using a AxioCam MRm camera and AxioVision software (Zeiss).

### Genetic mosaic analysis

To associate the somatic focus of action of *pod-2* for germline phenotypes, array losses were characterized in animals of genotype *pod-2(1691null); tnEx212[W09B6-pod-2(+)—20 ng/μl—hi copy; sur-5::gfp]* that were isolated on the basis of sterility. Subsequently, potential candidate mosaics of interest were prescreened directly at the dissecting microscope level for asymmetry in the phenotypic severity seen between anterior and posterior gonad arms, and then examined at the compound microscope level for loss of *sur-5::gfp* expression. Potential *pod-2(tn1691null); tnEx219[W09B6-pod-2(+)—4 ng/μl—lo copy; sur-5::gfp]* genetic mosaics were also prescreened in this manner. Intestinal (E–) mosaics were prescreened using low-power fluorescence SMZ18 Nikon (Melville, NY, USA) microscope with a Lumencor (Beaverton, OR, USA) SOLA Light Engine, but for *pod-2* many of these candidate mosaics were seen to express *sur-5::gfp* at very low levels, or in just a few intestinal cells, when examined at high power. This was not an issue when isolating *fasn-1(tn1762null); tnEx218[fasn-1(+); sur-5::gfp]* E(–) mosaics.

### Oligonucleotides used

#### *fasn-1(tn1762)* isolation

Single guide pairs:

5’-tcttggtatcttcccactcactga-3’ plus 5’-aaactcagtgagtgggaagatacc-3’, and 5’-cttggatgactttccttgcacga-3’ plus 5’-aaactcgtgcaaggaaagtcatcc-3’.

Repair template:5’- cgtatgacgtgatttttaatgtgataattactctgaaatatctcaaaacgtgcaaggaaagtcatcttcagtaacagtt gaacacatcaacaggattat-3’

Verification of *tn1762*:

5’- ctgacaagacgacggacaactcagg-3’ and 5’- ctacgtaaccgtaacggcatggcac-3’

#### *pod-2(tn1691)* isolation

Single guide pairs:

5’-tcttggaaaactgtccatatacaa-3’ plus 5’-aaacttgtatatggacagttttcc-3’, and 5’-tcttgacagaacaggaaaaagtgg-3’ plus 5’-aaacccagtttttcctgttctgtc-3’.

Repair template: 5’- ccgattttttgtgcaatttcagagcaatataagtatataaacgttattttcagagccatcactttttcctgttctgtgatata caagaaacctattcagtgccttatttgattgacgactac-3’

Verification of *tn1691*employed oligos:

5’-cgacgaagtaagtgagcctaattgtac-3’ with 5’-ttgtcaaccaccttacagtggcatg-3’

#### *gfp::pod-2(tn1765)* isolation

Single guide pair:

5’-tcttgcagggttacggatcttgga-3’ and 5’-aaactccaagatccgtaaccctgc-3

Homology arm oligo pairs:

5’-acgttgtaaaacgacggccagtcgccggcatctacctgtgtacctgacctgaccagattc-3’ plus

5’-aactccagtgaacaattcttctcctttactcattgtgaatgccagtccaagatccgtaac-3’;

5’-cgtgattacaaggatgacgatgacaagagagttgtaaacgggcaaaaaccagatatcag-3’ plus

5’-tcacacaggaaacagctatgatatgtgagagtgaatgaactgctccatgctctc-3’

#### *gfp::fasn-1(tn1782)* isolation

Single guide pair:

5’-tcttgaaggatcgtccgattcgag-3’ plus 5’-aaaactcgaatcggacgatccttc-3’

Primer used to introduce silent mutations to recognition site:

5’-gataatactggagagggctcatcagactcaagtggaacttgggagcgaatttcggac-3’

Homology flanking GFP:

5’-gggataataagtaggttctgacctcctctcccg-3’ to 5’-gatccattcgtgcttcttcttgtcccgtgag-3’

## Acknowledgements

We are grateful to our colleagues Gabriela Huelgas-Morales, Caroline Spike, Tatsuya Tsukamoto, and Jocelyn Shaw for discussions regarding experiments and comments on the manuscript. We would also like to thank Ross Johnson and David Hall for many years of sharing their knowledge and enthusiasm in all things concerning gap junctions. This work was supported by National Institutes of Health (NIH) grant GM57173 to DG. Some strains were provided by the Caenorhabditis Genetics Center, which is funded by grant P40OD010440 from the NIH Office of Research Infrastructure Programs.

## Additional information Funding

### Funding

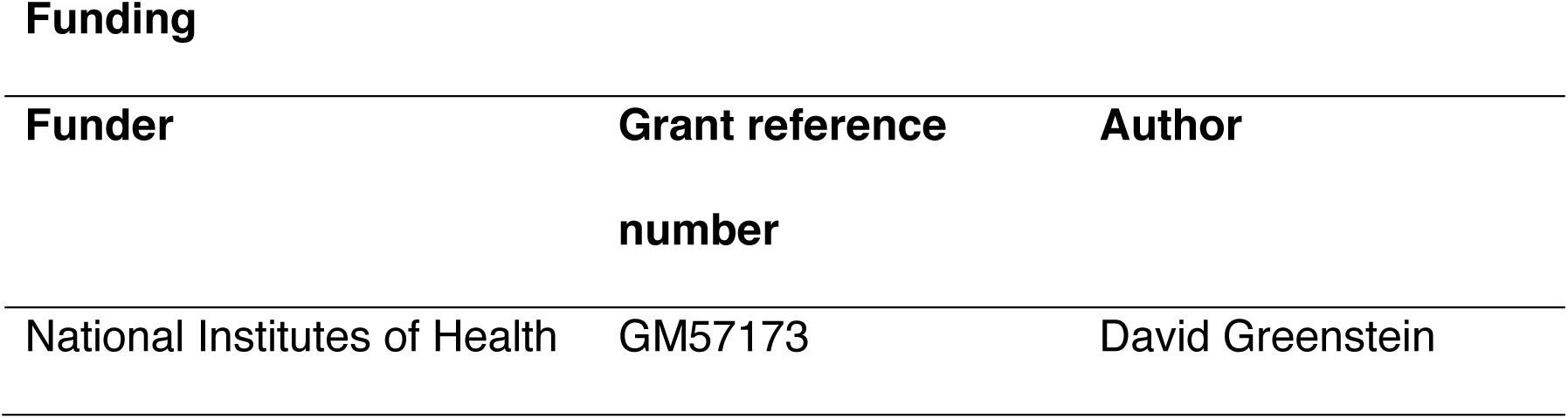

### Author contributions

TS conducted the experiments, prepared the figures, and wrote the manuscript. TS and DG designed and conceptualized the research study. DG obtained funding, supervised the study, and edited the manuscript.

## Figure Legends

**Figure 1–figure supplement table 1.**
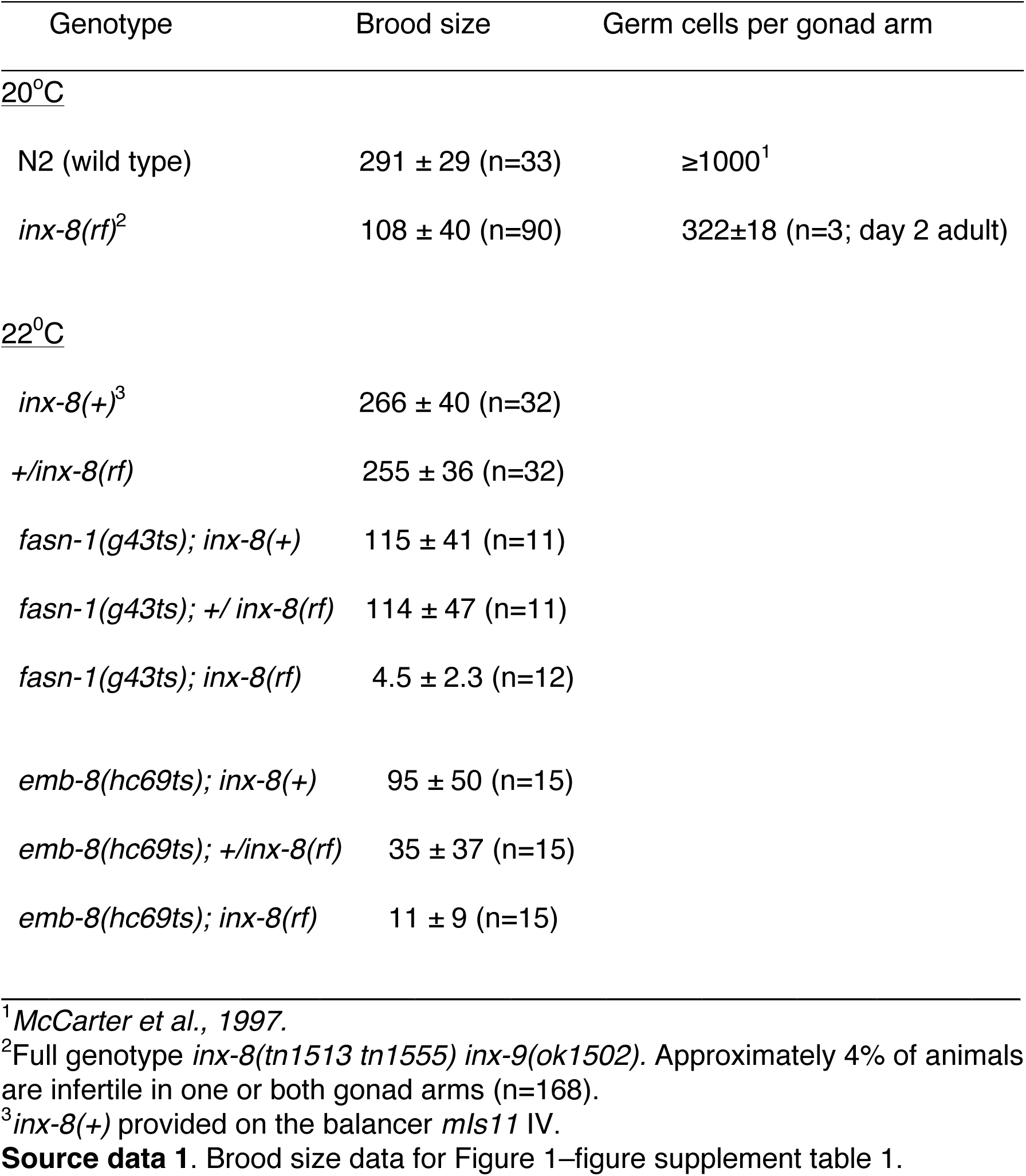
*inx-8(rf)* exhibits reduced brood size and interacts with FA synthesis pathway genes.

**Figure 1–figure supplement table 2.**
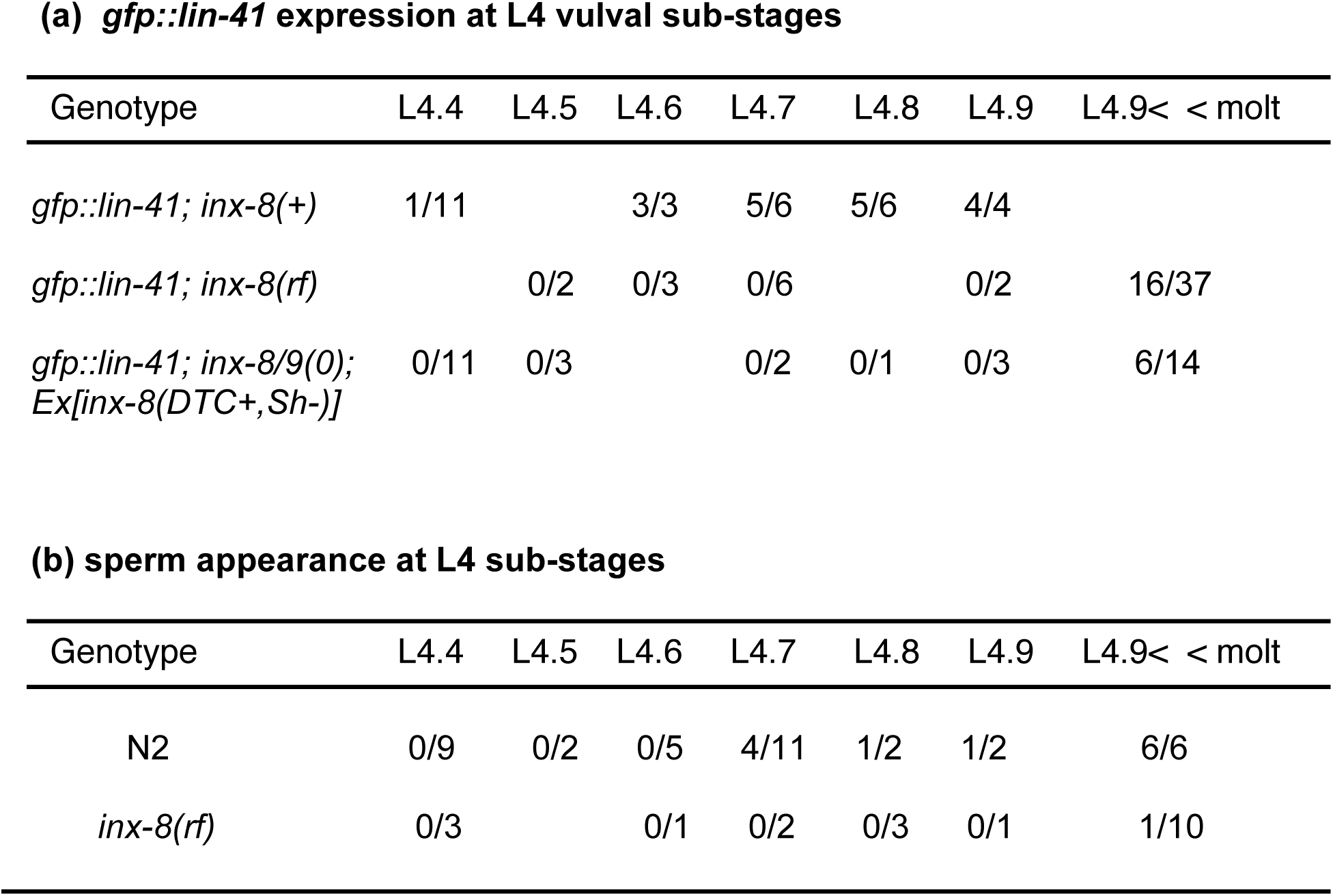
Reduced soma-germline gap junction coupling delays gametogenesis.

## Legends to Figure Supplements

**Figure 1–figure supplement 1.**
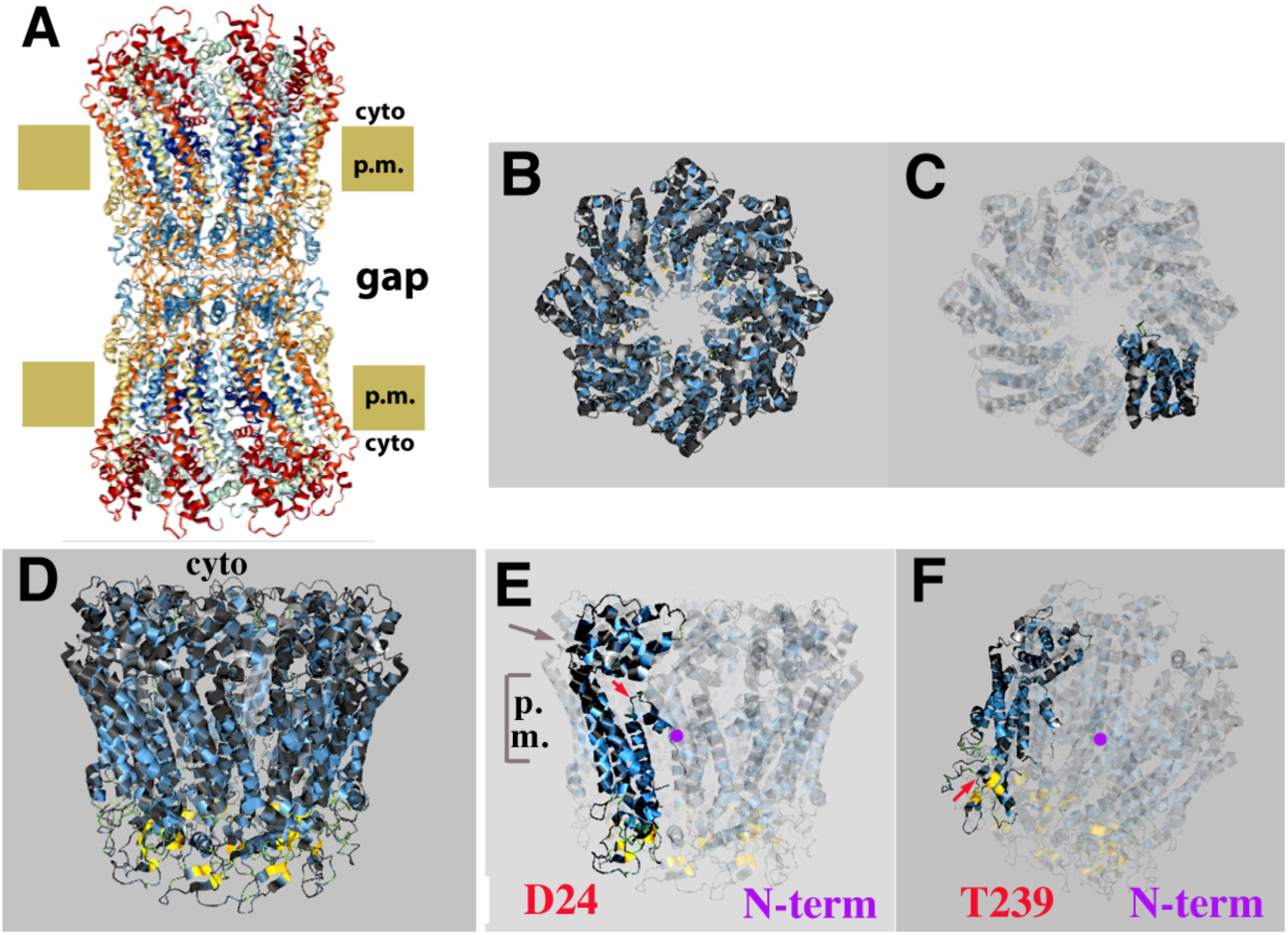
Predicted topological sites of amino acids mutated in *inx-8(tn1513 tn1555) inx-9(0)*. (A) Atomic structure of the *C. elegans* homomeric, homotypic INX-6 gap junction full channel, determined by *Oshima et al., (2016)* (PDB I.D. 5H1R, rcsb.org (*Berman et al., 2000*), viewed with NGL viewer (*Rose et al., 2018*). Octameric hemichannels associate via extracellular loops to form an enclosed channel in the region classically identified by TEM as the “gap” between closely aligned plasma membranes. cyto, cytoplasmic membrane face; p.m., plasma membrane. (B-F) Model for INX-8 hemichannel based on homology alignment to INX-6 (aquaria.ws). (B) Cytoplasmic view. (C) Single subunit highlighted. In wild type this hemichannel is comprised of both INX-8 and INX-9 subunits, and associates with germline hemichannels comprised of INX-14+INX-21 or INX-14+INX-22. (D) Side view of hemichannel. Beta-strands (yellow) in the extracellular loops contribute to walling off the pore. (E) Single subunit highlighted. Kinked region (gray arrow) allows the cytoplasmic structure of the subunits (cytoplasmic loop and C-terminus) to form a dome at the pore entry. Amino termini (N-term) line the pore as a funnel; a loop in the amino terminus allows for an interaction with residues in the dome (*Oshima et al., 2016*). D24 in INX-8 (red arrow) lies near the D21 residue predicted by homology with INX-6 (D25) to interact with the cytoplasmic dome. (F) INX-8 T239 is predicted to lie adjacent to a beta-strand contributing to hemichannel-hemichannel association (red arrow). (To expose the extracellular loop region in relation to (E), the aspect of view is rotated towards the viewer and angled up to the left.)

**Figure 2–figure supplement 1.**
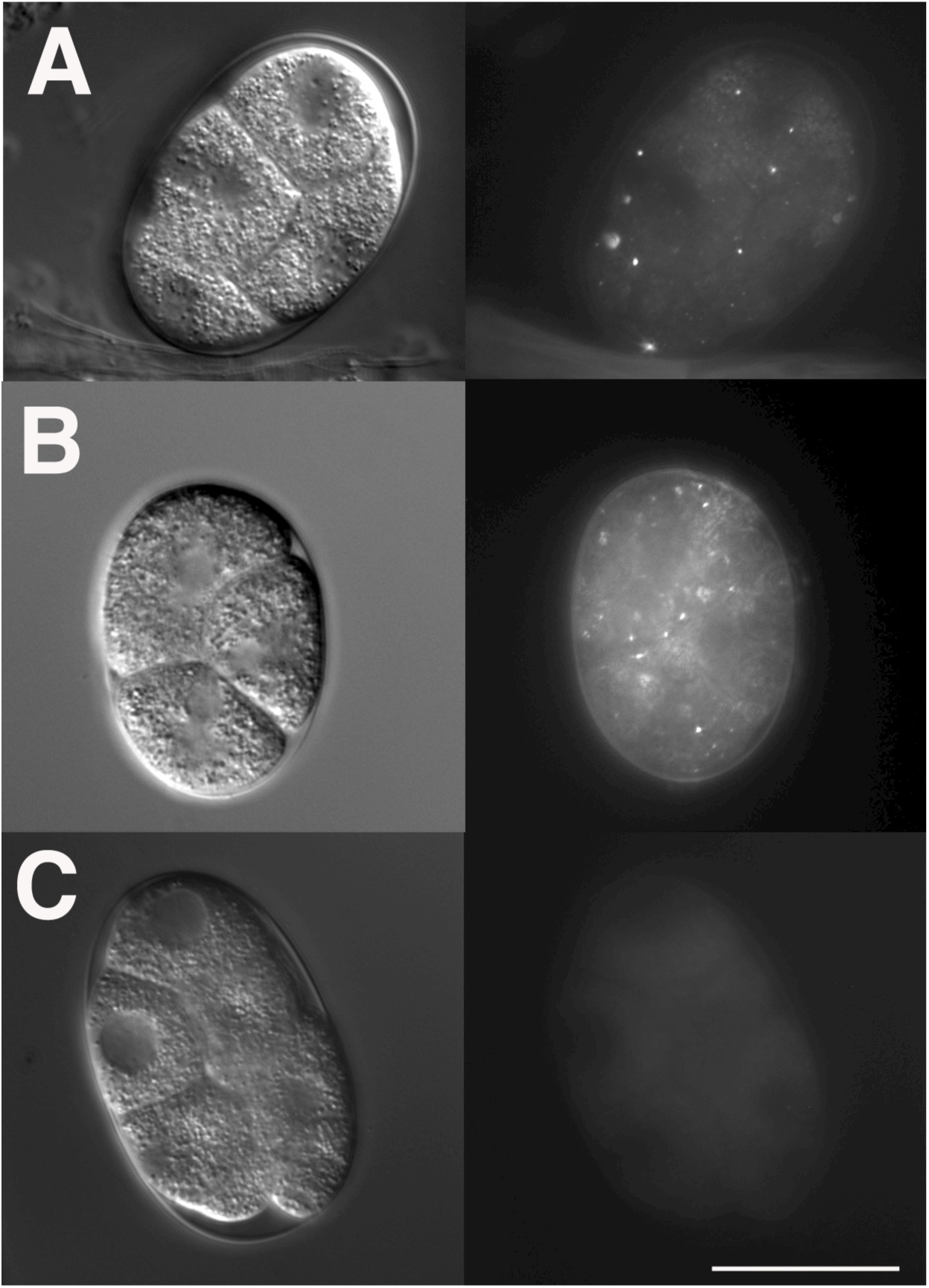
INX-8(rf)::GFP can be endocytosed by oocytes. (A) *inx-8(+)::gfp* expressed in the somatic gonad sheath is endocytosed during ovulation and appears as distinct fluorescent puncta in early embryos. (b) *inx-8(rf)::gfp* expressed on an extrachromosomal array in *inx-8(0) inx-9(0)* animals is endocytosed by ovulating oocytes. (c) *inx-8::gfp* expressed by the DTC in *inx-8/9(0); Ex[inx-8(DTC+,Sh-)]* is not observed in early embryos. Bar, 20 μm.

**Figure 7–figure supplement 1.**
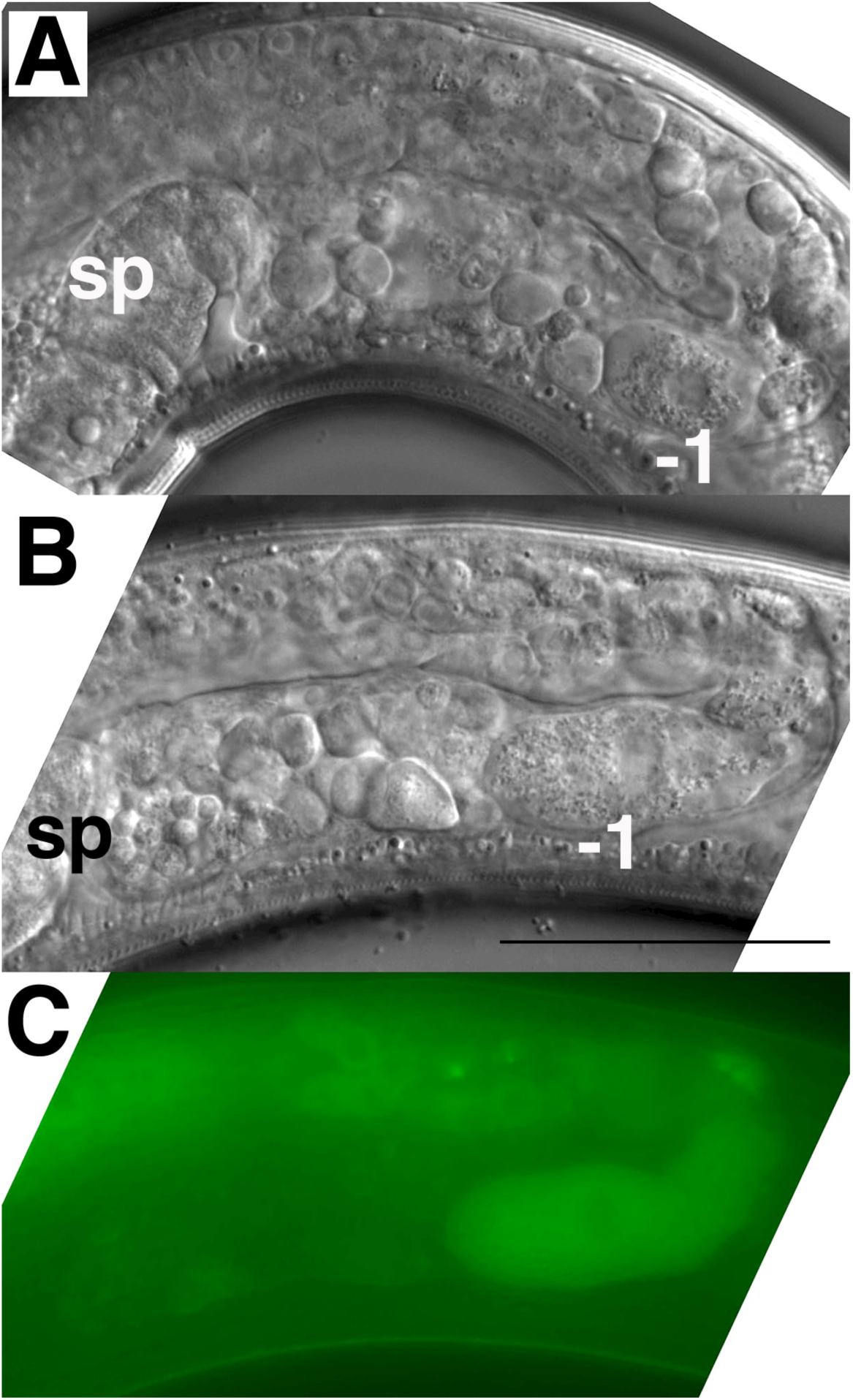
Expression of *gfp::pod-2* in the germline fails to rescue *inx-8(rf)* phenotypes. (**A**) Young (first-day) adult *inx-8(rf)* gonad arm. (B, C) Young adult *inx-8(rf)* gonad arm expressing *mex-5p::gfp::pod-2::tbb-2-3’UTR*. The *pod-2* isoform in this construct corresponds to *gfp::pod-2* isoform “a” in *Figure 3B*. This isoform encompasses sequence homology to all of the enzymatic sites in other ACC multi-functional enzymes. Sp, spermatheca; −1, most proximal oocyte; bar, 50 μm.

